# Instance-Wise Contrastive Graph Neural Network Enables the Discovery of Novel *Aedes aegypti* Larvicidal Compounds

**DOI:** 10.64898/2026.05.28.726277

**Authors:** Karine Santana Lúcio da Costa, Gustavo Henrique Garcia Caldeira, Vinícius Alexandre Fiaia Costa, Amanda Santana Silva, Carolina da Silva Pereira, Bruna Cassimiro Batista, Luana Bafutto Manchein, Holli-Joi Martin, Jamal Rafique, Rodolpho de Campos Braga, Eugene Muratov, Sumbal Saba, Gisele Augusto Rodrigues de Oliveira, Christian Luz, Juscelino Rodrigues, Bruno Junior Neves

## Abstract

*Aedes aegypti* remains a major arboviral vector, making larval control a critical strategy to reduce mosquito populations. However, resistance to commercial larvicides has reduced the long-term effectiveness of current interventions, reinforcing the need for new compounds with improved potency and selectivity. Here, we present an instance-wise contrastive graph neural network (GNN) framework to accelerate the discovery of novel larvicidal compounds. The model was trained on a curated dataset of 556 organic compounds organized into LC_50_-derived multitask classification thresholds and integrated Transformer-inspired graph learning with whole-molecule and fragment-level contrastive regularization. This model achieved strong held-out performance, with global AUC = 0.95 ± 0.01, PR-AUC = 0.93 ± 0.01, and MCC = 0.77 ± 0.03, outperforming conventional machine learning and graph-based baselines. Predictive uncertainty analysis and counterfactual maps further supported the interpretation of threshold-sensitive predictions and substructural contribution patterns. The model was applied to screen 1.3 million compounds, resulting in 10 candidates for experimental validation. Three compounds showed measurable larvicidal activity against *A. aegypti* larvae. Among them, **LC-79** emerged as the most promising hit, with 2-day and 5-day LC_50_ values of 0.24 µg/mL (0.66 µM) and 0.05 µg/mL (0.13 µM), respectively, an IE_50_ of 0.06 µg/mL (0.16 µM), and rapid larval mortality (LT_50_ = 1.10 days at 1 µg/mL). **LC-79** also showed no measurable acute toxicity to *Daphnia magna* at the highest tested concentration [EC_50-48h_ >43 µg/mL (>119 µM)], resulting in selectivity indices >180 and >860 relative to its 2-day and 5-day LC_50_ values. Overall, this study demonstrates that contrastive graph learning can move beyond retrospective larvicide modeling to experimentally validated hit discovery, identifying **LC-79** as a potent and preliminarily selective acylthiourea larvicide candidate for further mechanism-of-action, resistance, and semi-field evaluation.

## 1. Introduction

Vector-borne diseases remain among the most important infectious threats worldwide, accounting for more than 17% of all infectious diseases and causing more than 700,000 deaths annually.^1^ Among mosquito vectors, *Aedes aegypti* has major epidemiological relevance as an efficient transmitter of dengue,^2^ chikungunya,^3^ Zika,^4^ and yellow fever^5^ viruses in urban and periurban environments. The public-health impact of *Aedes*-borne diseases has intensified in recent years, with dengue reaching unprecedented global levels in 2024^6^ and expanding across regions historically considered less vulnerable.^7–9^ This trend is driven by the convergence of urbanization, climate change, climatic suitability, human mobility, and the ecological adaptation of *Aedes* mosquitoes to artificial breeding sites.^8,9^

Chemical larval control is a key component of integrated vector management, suppressing immature mosquito populations before adult emergence and pathogen transmission.^10,11^ Current interventions depend on a restricted set of active ingredients, including organophosphates, insect growth regulators, and spinosyns.^12,13^ However, the long-term effectiveness of interventions is constrained by the emergence of insecticide resistance.^14–18^ These limitations are particularly relevant for compounds deployed in domestic or peridomestic water containers, where repeated exposure can impose strong selection pressure on *A. aegypti* populations. In addition, their direct application to water-holding containers also requires careful consideration of potential effects on non-target organisms.^19–24^ Consequently, there is a pressing need for innovative discovery strategies to expand the chemical space of larvicidal agents for *A. aegypti* control.

Identifying new larvicidal hits remains a significant challenge, as conventional experimental screening is labor-intensive, resource-demanding, and poorly scalable for large chemical libraries. Artificial intelligence (AI)-driven approaches can accelerate larvicide discovery by prioritizing candidates with higher predicted activity for experimental validation.^25–27^ Within this context, machine learning has been the main strategy used in quantitative structure-activity relationship (QSAR) studies^28,29^ aimed at predicting larvicidal compounds against *A. aegypti*.^30–34^ These previous models demonstrated the feasibility of computationally modeling larvicidal activity, but their chemical coverage and generalizability remain limited to narrow chemotype-centered series. In addition, most of these models encode chemical structures through predefined molecular descriptors or fingerprints, which compress molecular topology into fixed feature vectors.^30–34^

Graph neural networks (GNNs) directly address this representational limitation by learning molecular representations directly from graphs. Formally, a molecular graph is defined as *G*= (*V, E*), in which *V* denotes the set of atoms and *E* denotes the set of bonds. Atom attributes are encoded in a feature matrix *X*, with each atom 𝒱 ∈ *V* represented by an initial vector *x*_𝒱_ ∈ ℝ^*d*^, whereas bond attributes are stored as edge features. Molecular connectivity is captured by an adjacency matrix *A*, where *A*_*ij*_ = 1 indicates a bond between atoms *𝒱*_*i*_ and *𝒱*_*j*_.^35,36^ Through graph-learning operations, atom embeddings are iteratively updated from neighboring atoms and bond features, allowing local functional groups, ring systems, and broader molecular context to be incorporated into the final representation.^37–39^ However, molecular activity may also depend on context-dependent relationships between distant or non-adjacent substructures. In this sense, recent advances in graph learning have incorporated Transformer-inspired attention mechanisms to complement message passing. These mechanisms adaptively weight atom- and bond-level interactions, improving the representation of long-range structural signals and chemically relevant substructural contexts.^40,41^

In this study, we developed an instance-wise contrastive GNN framework to identify novel larvicidal compounds against *A. aegypti*. This GNN architecture combines Transformer-inspired attention mechanisms with contrastive learning to integrate global- and local-chemical information within a unified molecular representation framework. The trained model was then applied to a sequential virtual screening campaign of 1.3 million commercially available compounds, followed by experimental validation in *A. aegypti* larvae and ecotoxicological evaluation using *Daphnia magna*. This workflow identified **LC-79** as a nanomolar larvicidal hit with rapid time-to-effect, adult-emergence inhibition, and low acute toxicity to non-target aquatic organisms.

## 2. Materials and Methods

### 2.1 Computational approaches

#### 2.1.1. Data collection and curation

A total of 707 chemical entries with reported larvicidal activity against *A. aegypti* were collected from the ChEMBL (target ID: CHEMBL613468)^42,43^ and ECOTOXicology knowledgebase.^44^ Entries with median lethal concentration (LC_50_) values determined against third- and fourth-instar larvae after 24 or 48 h of exposure were prioritized to ensure endpoint consistency. Since the reported LC_50_ values were available in different concentration units (e.g., mg/L, µg/mL, mg/mL, ppm, and ppb), all bioactivity data were standardized to micromolar units using molecular weight-based conversions when required. Subsequently, the LC_50_ values were transformed into a multitask binary classification matrix using four threshold-defined tasks: 0.1, 1, 10, and 100 µM. For each task, compounds with LC_50_ values ≤ the corresponding cutoff were labeled as active, whereas compounds with LC_50_ values > threshold were labeled as inactive. These cutoffs represent one-log-unit intervals and were adopted to provide progressively more stringent potency thresholds for subsequent predictive modeling analyses. Chemical structures in the Simplified Molecular-Input Line-Entry System (SMILES) format and corresponding bioactivity data were meticulously curated according to the guidelines proposed by Fourches *et al*.^45,46^ Curation steps included removing salts, mixtures, polymers, and organometallic compounds, and standardizing tautomeric forms to ensure dataset consistency. For duplicate entries with discordant activities, all corresponding records were excluded, while duplicates with concordant activities were consolidated to a single representative entry.

#### 2.1.2. Data description

The chemical space formed by actives and inactives was analyzed by plotting a similarity map generated with OSIRIS DataWarrior software v.05.02.01.3. The map was generated using the Rubberbanding Force Field approach, in which compounds are represented as nodes and pairwise chemical similarities define the forces that drive their spatial organization. The approach involves the following steps: (*i*) randomly positioning all compounds in 2D space; (*ii*) calculating the similarity matrix between all compounds using Tanimoto coefficients (Tc) and FragFP descriptors; (*iii*) identifying the most similar neighbors (Tanimoto coefficient >0.8) to be considered for each compound; and (*iv*) stepwise relocation of all compounds to ensure similar molecules were located close to each other.^47^

#### 2.1.3. Split of the dataset

The curated dataset was partitioned into training, validation, and test sets at an 8:1:1 ratio, using both random and scaffold-based splitting schemes. In the random split, compounds were assigned to subsets via uniform sampling. For the scaffold split, compounds were grouped by Bemis-Murcko scaffolds, ensuring each group contained only a single subset to prevent data leakage and enhance generalization assessment. This partitioning strategy was repeated five times using different random seeds, generating five independent train/validation/test splits for each splitting scheme. For each split, models were trained on the training set, hyperparameters were tuned on the validation set, and final performance was reported on the held-out test set to provide an unbiased estimate of predictive performance.

#### 2.1.4. Feature Representation

Molecular graphs were generated from SMILES using RDKit, with atoms represented as nodes and bonds as edges. Atom-level features included one-hot encoding of atomic number, degree, formal charge, hybridization state, and aromaticity. Bond-level features were encoded using one-hot representations of bond type, conjugation status, ring membership, and stereochemistry, including chirality. Feature vectors were built as tensors to provide structured graph representations during model training.

#### 2.1.5. Graph neural network

The proposed model was implemented as an atom-bond attentive message-passing neural network. At each message-passing layer, bond attributes were projected into a hidden space and processed using a FastFormer-inspired^48^ bond-attention mechanism. For a given molecular graph, bond embeddings were represented by the matrix *B* ∈ ℝ^*L*×*d*^, where *L* denotes the number of bonds and *d* denotes the hidden dimension. Bond-level projections were computed according to Equation 1:

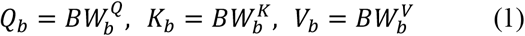

where *Q*_*b*_, *K*_*b*_, and *V*_*b*_ represent the bond-level query, key, and value matrices, respectively, computed independently for each attention head, and 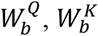, and 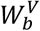 are learnable projection matrices. As shown in Equation 2, a feature-wise attention operation was applied to the bond-query representation to generate a global query vector:

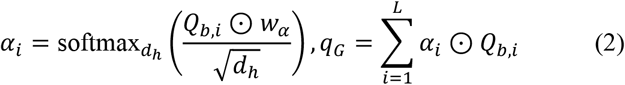

where *Q*_*b,i*_ denotes the query vector associated with bond *i, d*_*h*_ is the dimension of each attention head, *w*_*α*_ is a learnable attention vector, and ⊙ denotes element-wise multiplication, *α*_*i*_ is the corresponding feature-wise attention coefficient, *q*_*G*_ represents the global bond-query context aggregated over all valid bonds in the molecular graph, and *L* is the number of bonds. The notation softmax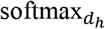 indicates that normalization is applied along the feature dimension of each attention head. The resulting global query vector was then used to modulate bond-key representations, as described in Equation 3:

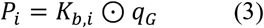

where *p*_*i*_ denotes the query-modulated key representation for this bond, and *K*_*b,i*_ is the key vector of bond *i*. A second feature-wise attention operation was subsequently applied to obtain a global key vector, as shown in Equation 4:

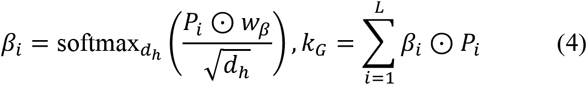

where *β*_*i*_ represents the feature-wise attention coefficient assigned to *p*_*i*_, *w*_*β*_ is a learnable attention vector, and *k*_*G*_denotes the global bond-key context obtained after aggregating the modulated key representations across bonds. Finally, the contextualized bond representation was calculated according to Equation 5:

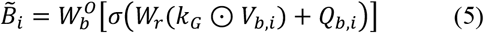

where 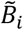 represents the final contextualized embedding of bond *i*, obtained by combining the bond-specific value vector *V*_*b,i*_ with the global bond-key context *k*_*G*_, followed by residual integration with the bond-query vector *Q*_*b,i*_. In addition, *W*_*r*_ and 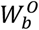 are learnable transformations, σ denotes the activation function, and ⊙ represents element-wise multiplication. The resulting contextualized bond embedding was subsequently used during message passing, enabling atom updates to incorporate information from both local bond features and the global bond environment. After message passing, atom embeddings were refined using a multi-head attention (MHA)^49^ readout block. The atom embeddings were represented by the matrix *H* ∈ ℝ^*N*×*d*^, where *N* denotes the number of atoms and *d* denotes the atom embedding dimension. Atom-level projections were computed according to Equation 6:

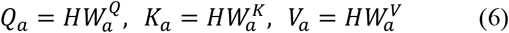

where *Q*_*a*_, *K*_*a*_, and *V*_*a*_ denote the atom-level query, key, and value matrices used to compute atom– atom attention, whereas 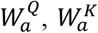, and 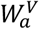 are learnable projection matrices. For each attention head, atom-level attention was calculated using structurally biased scaled dot-product attention, as defined in Equation 7:

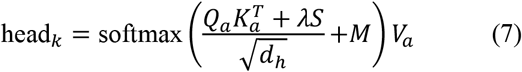

where head_*k*_ denotes the attention-weighted atom representation generated by the *k*-th attention head, *S* represents the structural bias matrix derived from molecular connectivity, *λ* is a scaling factor, and *M* is a graph mask that prevents attention between atoms belonging to different molecules in the same mini-batch, 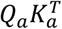 represents pairwise atom–atom compatibility scores, and *d*_*h*_ is the head-specific embedding dimension used for scaled dot-product attention. The structural bias incorporated random-walk positional encoding (RWPE), centrality-based encodings, and Laplacian-filtered positional information before computing atom–atom attention. The outputs from all attention heads were concatenated and linearly projected according to Equation 8:

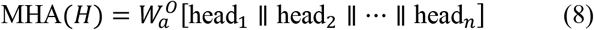

where MHA(*H*)represents the final atom-level attention output obtained after integrating the *n* head-specific representations into a single updated atom embedding matrix, and 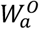 is the output projection matrix. The refined atom embeddings were finally aggregated into graph-level molecular embeddings and passed through fully connected layers to generate task-specific logits for larvicidal activity prediction.

#### 2.1.6. Composite loss function

Model optimization was performed using a composite loss function combining supervised multitask learning with molecular and fragment-level regularization. At the whole-molecule level, two augmented graph views were generated for each compound by random atom masking and random bond deletion. These paired views were treated as positive contrastive pairs, whereas augmented views from different compounds within the same batch were treated as negative pairs. The graph-level contrastive terms, inspired by the iMolCLR,^50^ were implemented as a weighted, temperature-scaled, normalized cross-entropy loss (NT-Xent)^51^ applied to projected molecular embeddings. In this formulation, SMILES-derived extended-connectivity fingerprints^52^ with diameter 4 (ECFP4) were used to compute pairwise Tanimoto similarity between compounds within each batch, and these similarity values were converted into negative-pair weights to reduce the contribution of structurally related molecules when they appeared as negatives. The weighted NT-Xent loss was calculated according to Equation 9:

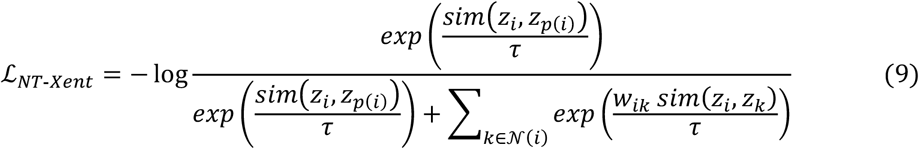

where *z*_*i*_ is the anchor embedding, *z*_*p*(*i*)_ is its positive augmented view, 𝒩 (*i*) denotes the set of negative examples in the batch, sim denotes cosine similarity, *τ* is the temperature parameter, and *w*_*ik*_ is the structural weighting factor assigned to each negative pair. The weighting factor was defined as *w*_*ik*_ = 1 − *λT*_*ik*_, where *T*_*ik*_ is the Tanimoto similarity calculated from molecular fingerprints, and *λ* controls the strength of structural downweighting. Thus, close structural analogs were not treated as equally strong negatives, while the contrastive representations themselves remained graph-derived embeddings.

At the local level, molecular fragments were generated using the breaking of retrosynthetically interesting chemical substructures (BRICS) procedure. Atom embeddings assigned to the same BRICS fragment were aggregated to produce fragment-level representations.^53^ The standard NT-Xent loss was then applied to projected BRICS fragment embeddings, corresponding to Equation 9 with *w*_*ik*_ = 1. Therefore, contrastive regularization was applied at two complementary levels: a fingerprint-weighted whole-molecule level that preserved global compound identity and a BRICS-based fragment level that regularized local substructural representations.

The supervised component was calculated using binary cross-entropy (BCE) with logits for each compound–task pair. To balance the contribution of the LC_50_-derived tasks during multitask learning, the supervised ℒ_*sup*_ was then weighted using homoscedastic uncertainty.^54^ For each task *t*, a trainable parameter *a*_*t*_ = log 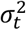 defined as the logarithm of the task-dependent variance was introduced to adaptively scale the ℒ_*sup*_ losses across tasks as shown in Equation 10:

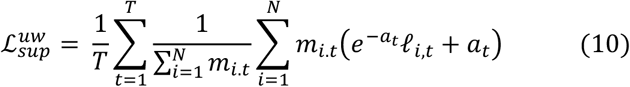

where 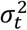 represents the homoscedastic uncertainty associated with task *t, T* is the number of tasks, *N* is the number of samples, *m*_*i,t*_ is a binary mask indicating whether compound *i* has a valid label for task *t*, and ℓ_*i,t*_ is the individual BCE loss calculated from the experimental binary label *y*_*i,t*_ and the corresponding model logit 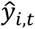. The exponential term *e*^−*a*_*t*_^ adaptively downweights tasks with higher uncertainty, while the additive *a*_*t*_ term penalizes overly large uncertainty estimates, preventing trivial solutions. The same uncertainty-weighting scheme was applied to the supervised losses calculated for the original molecular graph and for the augmented graph views. Thus, the augmented supervised term was defined as follows in Equation 11:

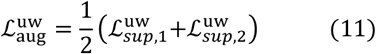

where 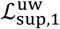 and 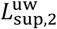 correspond to the uncertainty-weighted BCE losses calculated for the two augmented molecular views. The final training objective was then defined according to Equation 12:

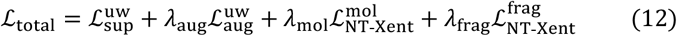

where 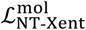 and 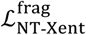 are the contrastive losses applied to whole-molecule and BRICS fragment-level embeddings, respectively. The coefficients *λ*_aug_, *λ*_mol_, and *λ*_frag_ control the relative contribution of supervised augmentation consistency, whole-molecule contrastive regularization, and fragment-level contrastive regularization.

#### 2.1.7. Monte Carlo dropout

Predictive uncertainty was estimated using Monte Carlo dropout during inference.^55,56^ The trained model was kept in evaluation mode, while dropout layers were reactivated to generate 30 stochastic forward passes for each compound. For each task, stochastic logits were converted to probabilities using the sigmoid function, and the resulting probability distributions were used to calculate uncertainty scores. For a given compound *i* and task *t*, normalized predictive entropy was calculated according to Equation 13:

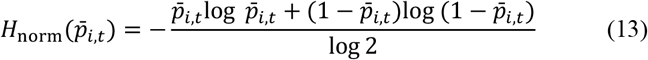

where 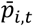 is the mean predicted probability for compound *i* and task *t* across stochastic forward passes. The normalization by log 2 bounds the entropy between 0 and 1 for binary predictions. Mutual information was used as a proxy for epistemic uncertainty and was calculated as the difference between predictive entropy and expected entropy across Monte Carlo

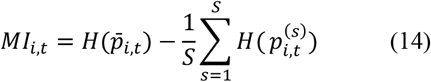

where *MI*_*i,t*_ denotes the mutual information for compound *i* and task *t, S* is the number of stochastic forward passes, 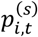 is the predicted probability obtained at stochastic sample *s*, 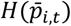 is the binary entropy of the mean predicted probability, and 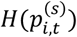 is the binary entropy calculated for each stochastic prediction. Here, *H*(*p*) denotes the non-normalized binary entropy, defined as −[*p*log *p* +(1 − *p*)log(1 − *p*)]. Finally, decision stability was assessed using the confidence margin, defined as the normalized distance between the mean predicted probability and the task-specific classification threshold *τ*_*t*_, according to Equation 15:

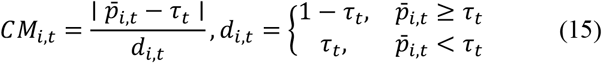

where *CM*_*i,t*_ represents the confidence margin and *τ*_*t*_ is the task-specific decision threshold. For predictions assigned to the active class, the denominator *d*_*i,t*_ corresponds to the maximum possible distance above the threshold, whereas for predictions assigned to the inactive class, it corresponds to the maximum possible distance below the threshold.

#### 2.1.8. Model training

The model was implemented in PyTorch v.2.5.1 with PyTorch Geometric v.2.6.1 and trained on an NVIDIA TITAN Xp GPU. Training proceeded for up to 500 epochs with a batch size of 24, employing early stopping to prevent overfitting. Training was stopped when the validation loss failed to improve within an adaptive five-epoch window; otherwise, the best validation loss checkpoint was refreshed. Regularization comprised gradient clipping and a cosine annealing learning rate schedule with a warm-up period. During hyperparameter search, pruning was enabled to terminate unpromising trials. Hyperparameters were tuned using Optuna v.4.2.0, conducting 100 trials to systematically explore the search space. Finally, embedding vectors were saved during training and projected onto a t-SNE map to monitor the organization of the latent space.

#### 2.1.9. Benchmarking

The proposed model was benchmarked against conventional machine learning and GNN baselines. Random Forest (RF) and Support Vector Machine (SVM) models were implemented in Scikit-learn v0.24.2 using ECFP4 fingerprints as input descriptors, with hyperparameters optimized by Bayesian optimization in Scikit-Optimize v0.7.4 under a 5-fold cross-validation scheme. Graph-based baselines, including Graph Attention Network (GAT),^58^ Attentive Fingerprint (AttentiveFP),^59^ Message Passing Neural Network (MPNN),^60^ and Graph Isomorphism Network (GIN),^61^ were implemented in PyTorch v2.5.1 with PyTorch Geometric v2.6.1. All models were trained and evaluated using identical data split protocols to ensure direct methodological comparability.

#### 2.1.10. Statistical analysis

Model performance was evaluated using accuracy (ACC), recall, specificity (SP), positive predictive value (PPV), negative predictive value (NPV), F1 score, Matthews correlation coefficient (MCC), and area under the curve (AUC). Additionally, the geometric mean (G-mean) and the precision-recall area under the curve (PR-AUC) were used to address the imbalance in the tasks. All metrics are reported as the mean ± standard deviation (SD) across five independent data splits. Model performances were analyzed using GraphPad Prism v8.0.2 (GraphPad Software, San Diego, CA, USA). Two-group comparisons were performed using an unpaired Student’s t-test or the Mann–Whitney U test, as appropriate.

#### 2.1.11. Model explainability

Model explainability was assessed using a counterfactual perturbation strategy adapted from Fiaia Costa *et al*.^62^ Briefly, selected compounds were systematically perturbed at local functional-group, bond, or ring environments using chemically valid isosteric transformations. For each valid perturbation, the change in predicted probability relative to the unperturbed molecule was calculated and assigned to the modified atomic environment. The resulting atom-level contribution scores were projected onto the molecular structure to generate contribution maps, highlighting substructural regions with positive or negative influence on the predicted larvicidal activity. Additionally, atom-level attention weights from FastFormer and MHA were extracted, providing a complementary visualization of substructural relevance.

### 2.2. Experimental approaches

#### 2.2.1. Chemicals

All compounds were purchased from ChemBridge (San Diego, CA, USA), dissolved in 1 mL of 10% (v/v) dimethyl sulfoxide (DMSO), and stored at -80 °C in 1.5 mL microtubes. Compound aliquots were thawed at room temperature and adjusted to 1 mL with 800 µL of 10% DMSO.

#### 2.2.2. Breeding and preparing adult mosquitoes

The *A. aegypti* colony, which originated from larvae collected in Goiânia, Brazil, in 2012, was maintained in the laboratory at 27 ± 5 °C and 75 ± 10% relative humidity under a natural photoperiod, as described by Rodrigues *et al*.^63^ Adults were fed *ad libitum* on filter paper imbibed with 10% sucrose solution, and females were fed twice a week following the method described by Lima *et al*.^64^ This procedure had been previously approved by the Ethics Commission for the Use of Animals of the Universidade Federal de Goiás, Goiânia (CEUA 050/20, UFG, 2020). Eggs laid on filter paper were transferred to dechlorinated tap water at 36 °C, and eclosed larvae were fed with ground cat food (Bom Preço^®^, Salto de Pirapora, Brazil). Second-instar larvae were transferred to distilled water and reared until molting. Newly molted third-instar larvae were starved for 12–24 hours before testing.

#### 2.2.3. Larvicidal bioassay

The larvicidal activity of selected compounds was evaluated following the World Health Organization guidelines for laboratory testing of mosquito larvicides, with some modifications.^65^ Five concentrations were prepared for each compound: 10, 1, 0.1, 0.01, and 0.001 µg/mL. Two positive controls, pyriproxyfen (PPF) and temephos (TEM), were included and tested at the same concentrations as the selected compounds. The negative control consisted of 1% (v/v) DMSO. Ten third-instar larvae were transferred to 25 mL of each solution and incubated at 25 ± 1 °C and 12 hours photoperiod (L:D 12:12). Larvae were fed 24 hours after the start of the bioassay and thereafter every 2 days. Larval mortality was recorded daily for up to five days after exposure. The surviving larvae were monitored for developmental progression until pupation and adult emergence for up to 15 days after exposure. The number of pupae and emerging adults was recorded to assess potential inhibitory or delayed effects on mosquito development. The LC_50_ was calculated at 2 and 5 days after exposure from larval mortality data, whereas the median emergence-inhibition concentration (IE_50_) was calculated after 15 days from adult-emergence inhibition data. Both endpoints were estimated by nonlinear regression using a normalized variable-slope concentration–response model in GraphPad Prism v8.0.2 (GraphPad Software, San Diego, CA, USA). The median lethal time (LT_50_), defined as the time required to induce 50% larval mortality at a fixed concentration, was estimated by probit analysis in Wolfram Mathematica 11.3 (Wolfram Research, Inc., Champaign, IL, USA).

#### 2.2.4. Ecotoxicological test

*D. magna* microcrustaceans were cultivated and maintained in a climatic chamber (SL-224, Solab) at 20 ± 2 ºC and 16 hours light:8 hours dark photoperiod. The culture medium was composed of deionized water with CaCl_2_ (11.76 g/L), MgSO_4_ (9.73 g/L), NaHCO3 (2.59 g/L), and KCl (0.23 g/L). Total hardness ranged from 175 to 200 mg CaCO_3_/L, with a pH of 7.0 and dissolved oxygen greater than 5 mg/L. The water was renewed weekly, and the organisms were kept in batches of up to 12 adults in 250 mL beakers. The culture was fed with algae of the species *Raphidocelis subcapitata* and with a commercial fish feed, Tetramin® (0.1 g/L), solubilized in ultrapure water. The culture medium was renewed three times a week. Neonates aged 6-24 hours, obtained by parthenogenesis from the second brood of females aged 7-28 days, were employed in the bioassays.

##### 2.2.4.1. Daphnia magna acute toxicity assay

Acute toxicity tests were conducted with *D. magna* following Organization for Economic Co-operation and Development (OECD) test No. 202.^66^ *D. magna* neonates were exposed to culture medium as the negative control, zinc sulfate heptahydrate at 10 mg/L as the positive control, and test compounds (**LC-79** and **LC-83**) at five serial concentrations: 0.00435, 0.0435, 0.435, 4.35, and 43 µg/mL. Neonates were randomly distributed into glass tubes containing 10 mL of the test solution, with five organisms per tube and four tubes per treatment (four replicates; n = 5 per replicate). The tubes were transferred to an incubator at 20 ± 2 ºC, with a 16 h photoperiod and without feeding for 48 hours. Immobility was estimated after 48 hours of exposure and was defined as organisms that could not swim within 15 seconds after the glass tube was gently agitated. The median effective concentration for immobilization (EC_50-48h_) was estimated by nonlinear regression using a normalized variable-slope concentration–response model in GraphPad Prism v8.0.2 (GraphPad Software, San Diego, CA, USA).

## 3. Results and Discussion

From an applied vector-control perspective, the key challenge is not only to predict whether a compound is active, but to prioritize compounds that are chemically distinct from heavily used larvicidal classes and sufficiently potent to justify experimental follow-up. This requirement is particularly important for *A. aegypti* control because operational larvicides are drawn from a limited number of chemical and mechanistic classes, and repeated exposure in domestic water containers can favor resistance selection. Therefore, the modeling objective in this study was designed to support prospective hit discovery rather than retrospective classification alone: the framework needed to preserve potency-dependent structure–activity information, remain useful under scaffold-level generalization, and enrich large commercial libraries for experimentally testable candidates.

### 3.1. Data description

A total of 707 chemical entries with reported larvicidal activity against *A. aegypti* were collected from ChEMBL,^42,43^ and ECOTOXicology knowledgebase.^44^ After standardization and structure curation, the final dataset comprised 556 unique compounds. Subsequently, compounds were categorized as active or inactive using 0.1, 1, 10, and 100 μM thresholds (Figure 1a). The stringent 0.1 μM threshold was applied to capture highly potent larvicidal profiles, resulting in 74 actives and 482 inactives. The 1 μM and 10 μM thresholds yielded 171 actives/385 inactives and 229 actives/327 inactives, respectively, whereas the 100 μM cutoff generated the most balanced classification setting, with 295 actives and 261 inactives. This threshold-based organization enabled the analysis of larvicidal activity patterns across distinct potency ranges.

**Figure 1.**
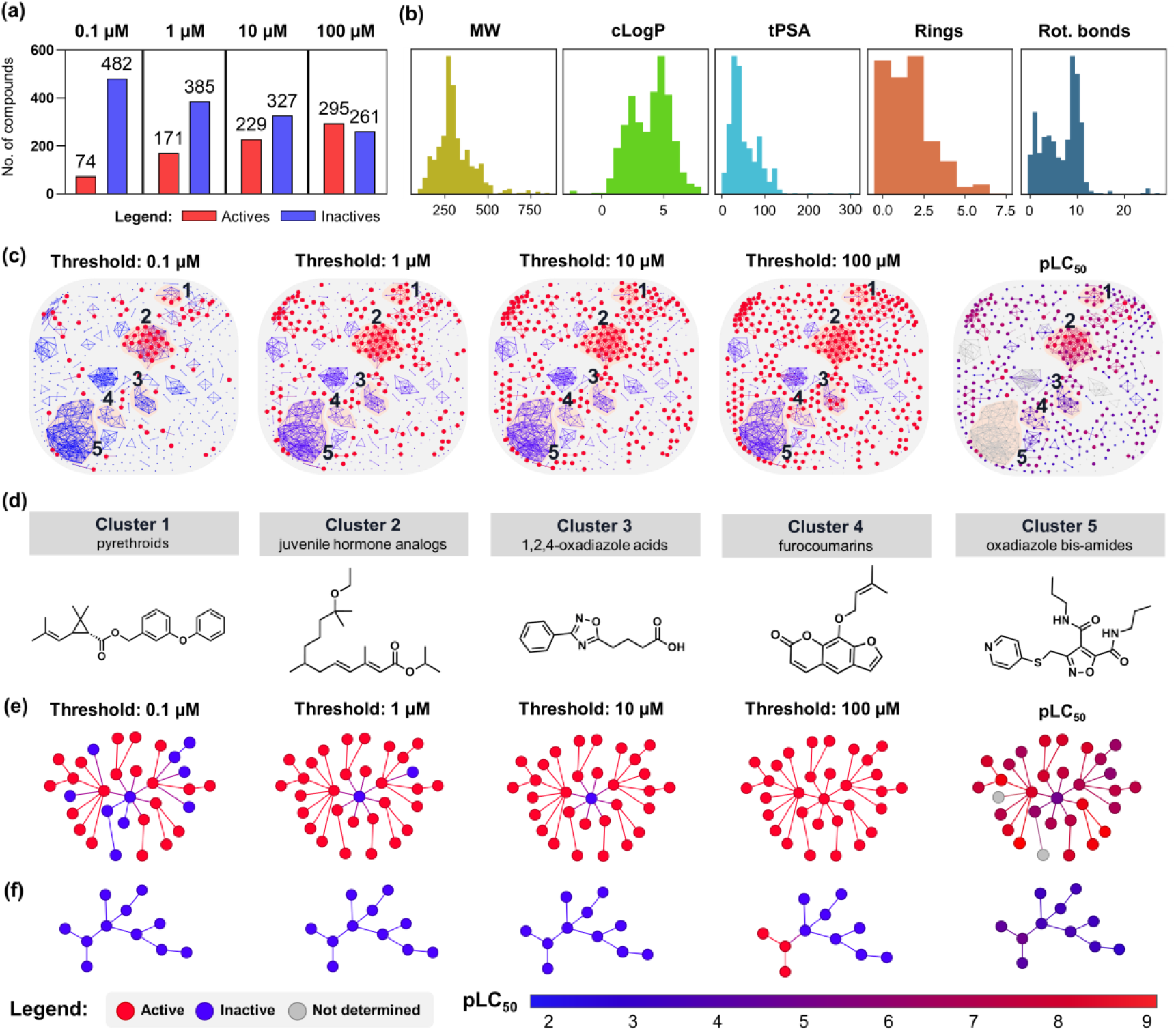
Chemical space analysis and structural profiling of active and inactive compounds in the dataset. (**a**) Distribution of actives and inactives across four LC_50_ classification thresholds. (**b**) Physicochemical space of the dataset is defined by MW, cLogP, tPSA, ring count, and rotatable-bond count. (**c**) Similarity maps showing the organization of actives and inactives across the four thresholds, along with the corresponding continuous pLC_50_ activity landscape. (**d**) Representative compounds from the main structural clusters identified in the dataset. (**e**) Isolated similarity subnetwork of cluster 2, comprising juvenile hormone analogs, extracted from the threshold-specific similarity maps. (**f**) Isolated similarity subnetwork of cluster 4, comprising furocoumarin derivatives. Red and blue nodes indicate active and inactive compounds, respectively; gray nodes indicate compounds without determined pLC_50_ data. pLC_50_ values are shown using a blue-to-red gradient.

The physicochemical profile of the curated dataset was examined using molecular weight, cLogP, tPSA, ring count, and rotatable-bond count (Figure 1b). The descriptor distributions indicated that the dataset is mainly populated by compounds with low-to-moderate ring complexity, with a large fraction containing 0–2 rings, and by structures with approximately 9–10 rotatable bonds. Chemical similarity maps further indicated that the dataset is organized into dominant structural clusters, in which actives and inactives showed partially overlapping, threshold-dependent distributions (Figure 1c). The continuous pLC_50_ landscape supported this pattern, revealing potency gradients within specific regions of the map rather than a homogeneous distribution of larvicidal activity.

Inspection of the similarity maps revealed five major structural clusters (Figure 1d) represented by pyrethroids, juvenile hormone analogs, 1,2,4-oxadiazole acids, furocoumarins, and oxadiazole-bis-amides. Juvenile hormone analogs (Figure 1e) and furocoumarin (Figure 1f) clusters were further examined as isolated subnetworks to illustrate how activity patterns change across distinct potency ranges. These clusters show that the composition of actives and inactives within the same chemotype is threshold-dependent, suggesting that a single activity cutoff may alter the apparent structural coverage of the active class. Such threshold-dependent organization indicates that larvicidal potency is not uniformly distributed across chemical space, supporting the use of a multitask learning strategy capable of preserving potency-dependent structural information.

### 3.2. Model performance

Accurate prediction of larvicidal activity requires molecular representations that capture both substructural patterns and broader graph-level context. This is particularly relevant in the present dataset because structurally related chemotypes may be assigned to different classes depending on the LC_50_ threshold. In addition, the most stringent cutoff results in a markedly imbalanced task, with only a limited number of highly potent compounds classified as actives. Under these conditions, models relying on fixed descriptors or conventional message passing may struggle to capture subtle substructural signals while integrating long-range molecular information. Therefore, we designed an instance-wise contrastive GNN framework to improve representation stability across related potency thresholds and to encode larvicidal activity at both whole-molecule and fragment levels.

The proposed framework integrates two contrastive learning components. At the whole-molecule level (Figure 2a), augmented molecular graphs derived from the same compound were treated as positive pairs, encouraging the model to preserve compound identity under chemically mild perturbations. At the fragment level (Figure 2b), BRICS-derived substructures were used to constrain the representation of local chemical environments, allowing the model to capture recurring substructural patterns associated with larvicidal potency. This dual contrastive strategy was designed to regularize graph embeddings while retaining both global molecular context and fragment-level chemical information.

**Figure 2.**
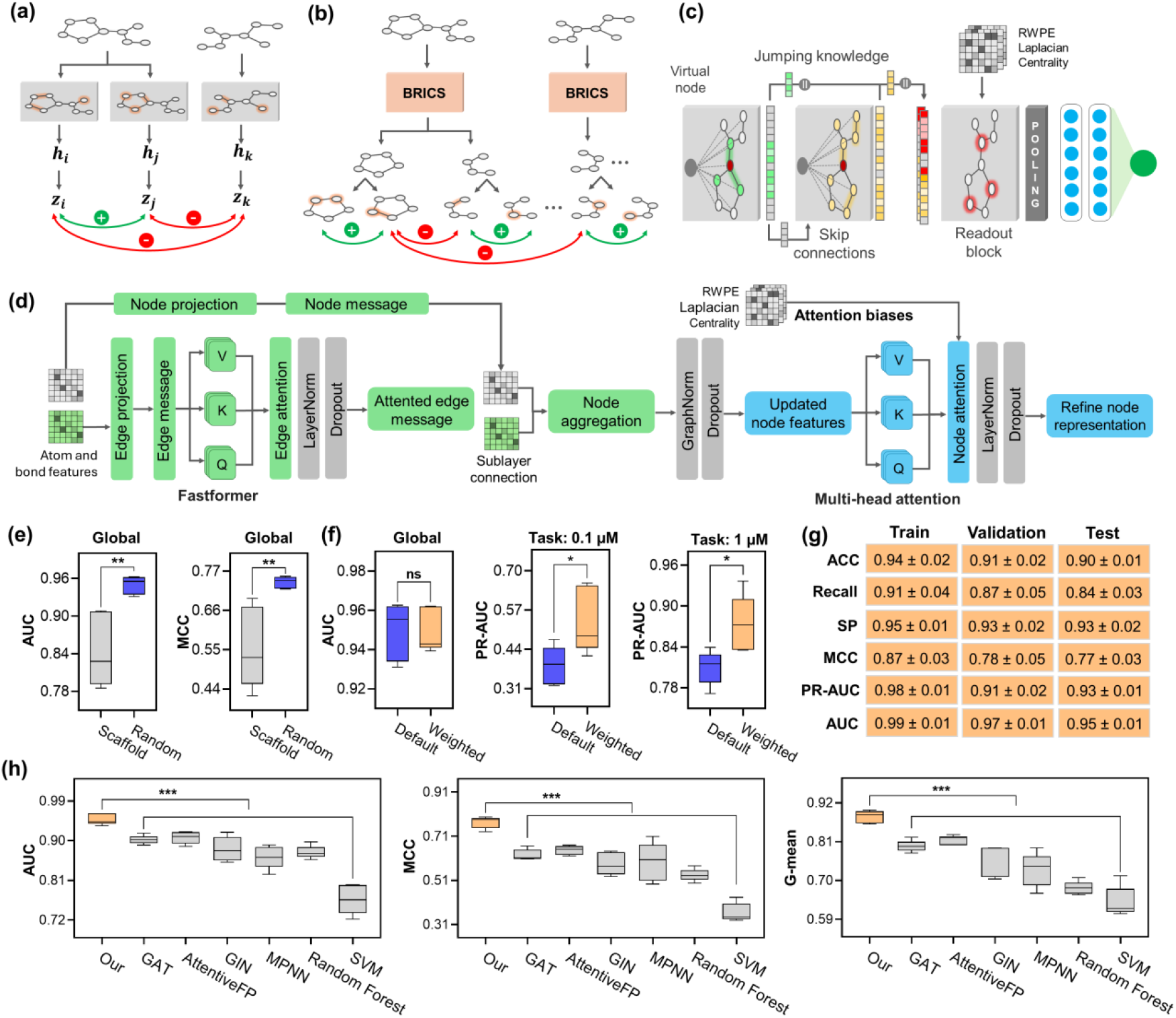
Instance-wise contrastive GNN design and predictive performance across split, weighting, and benchmarking analyses. (**a**) Whole-molecule instance-wise contrastive learning using augmented molecular graphs as positive pairs and unrelated compounds as negative pairs. (**b**) BRICS-based fragment-level contrastive learning, capturing local substructural representations across molecular fragments. (**c**) Overall architecture of the proposed GNN, integrating virtual nodes, skip connections, jumping knowledge, structurally informed readout, pooling, and task-specific prediction layers. (**d**) Detailed representation of the atom-bond message-passing workflow, featuring FastFormer-inspired edge attention and structurally biased MHA atom readout. (**e**) Comparison of random and scaffold splits based on global AUC and MCC. (**f**) Impact of homoscedastic uncertainty weighting on global AUC and task-specific PR-AUCs at 0.1 and 1 µM thresholds. (**g**) Global training, validation, and test set performances of the proposed model. (**h**) Benchmarking of the proposed model against GAT, AttentiveFP, GIN, MPNN, Random Forest, and SVM using AUC, MCC, and G-mean. * *p* < 0.05; **, *p* < 0.01; ***, *p* < 0.001, ns: not significant.

The GNN architecture combined atom-bond message passing with multiscale graph representation learning (Figure 2c). Virtual nodes, skip connections, and jumping knowledge were incorporated to enhance long-range information flow, retain signals captured at different message-passing depths, and mitigate oversmoothing. As shown in Figure 2d, bond features were processed using a FastFormer-inspired edge-attention module,^48^ as bond states determine how chemical information is propagated between neighboring atoms during message passing. This mechanism contextualizes each bond message against the global bond environment before node aggregation, improving the representation of chemically relevant linkers, ring bonds, and functional-group connectivity while maintaining computational efficiency. After message passing, atom embeddings were refined using a structurally biased MHA readout (Figure 2d).^67^ This second attention stage allowed the model to reweight atom-level environments in the final molecular representation, while RWPE, Laplacian features, and centrality descriptors provided topology-aware biases for capturing broader graph organization.

Model performance was first compared under random and scaffold split strategies (Figure 2e). The random split yielded a higher global test set AUC (0.95 ± 0.01) than the scaffold split (0.85 ± 0.06), with a statistically significant difference (*p* < 0.01). A similar pattern was observed for MCC, with random split performance (0.74 ± 0.02) exceeding scaffold split performance (0.56 ± 0.11, *p* < 0.01). Complete task-specific performances under random and scaffold split strategies are provided in Tables S1 and S2, respectively. The lower performance under scaffold splitting is consistent with the dataset organization described above, in which a limited number of dominant chemotypes contribute substantially to the chemical space. Under this condition, scaffold partitioning imposes a stricter generalization regime by separating structurally related compounds between the training and test sets.

Importantly, the scaffold-split performance should be interpreted as the more conservative estimate of prospective chemical-space extrapolation. The reduced MCC under scaffold splitting indicates that prediction of highly novel scaffolds remains more challenging than interpolation among related analogs. For this reason, the model was not used as a stand-alone decision engine; instead, predictions were combined with uncertainty assessment, chemical-similarity filtering, commercial availability, and experimental larvicidal validation. This framing is essential because the practical value of the workflow depends on enrichment of testable hits rather than perfect prospective classification across all possible chemotypes.

Due to differing distributions of actives and inactives across LC_50_-derived tasks, the instance-wise contrastive GNN was trained using homoscedastic uncertainty weighting in supervised losses. This approach adaptively scales task-specific losses through learned uncertainty parameters, reducing the dominance of individual tasks during training. As shown in Figure 2f, uncertainty weighting produced global AUC values close to the default supervised loss configuration (default: 0.95 ± 0.01; weighted: 0.95 ± 0.01), with no significant differences. However, task-specific PR-AUCs improved after uncertainty weighting for the most stringent thresholds. For the 0.1 µM task, PR-AUC increased from 0.39 ± 0.06 to 0.54 ± 0.11 (*p* < 0.05). A similar improvement was observed for the 1 µM task, increasing from 0.81 ± 0.02 to 0.87 ± 0.04 (*p* < 0.05). These results indicate that adaptive task weighting was particularly beneficial for potency thresholds with stronger class imbalance. The final weighted model maintained strong held-out performance across several indicators (Figure 2g), achieving ACC = 0.90 ± 0.01, recall = 0.84 ± 0.03, SP = 0.93 ± 0.02, MCC = 0.77 ± 0.03, PR-AUC = 0.93 ± 0.01, and AUC = 0.95 ± 0.01. The full task-wise performance profile of the uncertainty-weighted model is reported in Table S3.

The proposed model was further benchmarked against conventional machine learning and graph-based baselines using global test-set performances. Detailed task-specific performances for the baseline models are provided in Tables S4–S9. As shown in Figure 2h, it achieved statistically superior performance over all baseline methods across the benchmark comparisons (*p* < 0.001). For AUC, AttentiveFP (–4%) and GAT (–5%) were the closest graph-based baselines, followed by GIN (–7%) and MPNN (–9%). The conventional ECFP4-based models showed lower discriminative AUC performance, with Random Forest (–8%) and SVM (–18%) trailing the proposed model. The performance gap was markedly greater for metrics that are more sensitive to class imbalance. For MCC, AttentiveFP (–13%) and GAT (–14%) remained the strongest baselines, whereas GIN (–18%), MPNN (–21%), Random Forest (–24%), and SVM (–41%) showed greater reductions. A similar trend was observed for G-mean, with smaller decreases for AttentiveFP (–7%) and GAT (–8%), followed by GIN (–12%), MPNN (–15%), Random Forest (– 20%), and SVM (–24%).

### 3.3. Predictive uncertainty analysis

Predictive uncertainty was analyzed to determine how reliably the model separated active and inactive compounds across the LC_50_-derived tasks. As shown in Figure 3a, most classified compounds were found in regions of low mutual information and high confidence margin, indicating reliable predictions away from the decision boundary. This pattern was especially evident for the 0.1 µM task, in which the large inactive class was predicted with high confidence. However, the few active compounds at this stringent threshold were concentrated closer to low-margin regions, indicating that uncertainty was not uniformly distributed but was enriched among highly potent larvicides. The 1 and 10 µM tasks showed broader dispersion toward higher mutual information and lower confidence margins, suggesting greater decision-boundary uncertainty. In contrast, the 100 µM task showed a more stable classification profile, consistent with a broader, more populous active class. The normalized entropy distributions confirmed this class-dependent uncertainty pattern (Figure 3b). Across all tasks, inactives were concentrated at low entropy, whereas active compounds displayed broader entropy profiles. This effect was strongest for the 0.1 µM task, indicating that highly potent larvicides were more difficult to model because they constitute a small and chemically restricted class. At the less stringent thresholds, the broader entropy distributions among actives likely reflected greater structural and potency heterogeneity within the active class.

**Figure 3.**
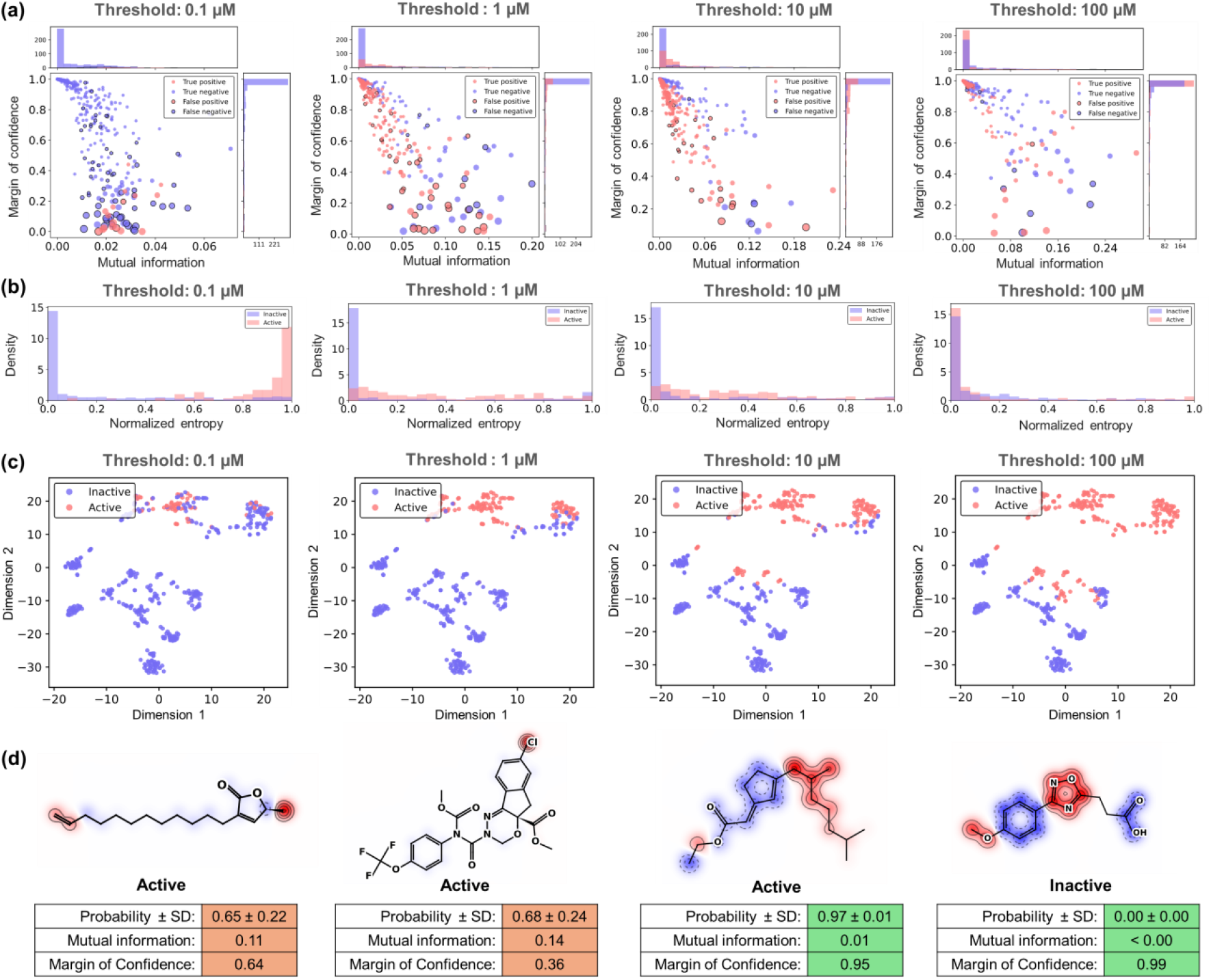
Predictive uncertainty of the instance-wise contrastive GNN model. (**a**) Mutual information versus confidence margin across the 0.1, 1, 10, and 100 µM LC_50_-derived tasks. (**b**) Normalized entropy distributions for actives and inactives across each task. (**c**) t-SNE projections of the learned molecular embeddings at the optimal training epoch, showing threshold-dependent organization of active and inactive compounds. (**d**) Representative counterfactual maps generated for the 10 µM task. Atom-level contribution maps are accompanied by predicted probability ± SD, mutual information, and confidence margin. Red and blue regions indicate positive and negative contributions to the predicted larvicidal activity, respectively.

The class-dependent uncertainty was further supported by the t-SNE projections of the learned molecular embeddings (Figure 3c). The most stringent 0.1 µM threshold yielded a sparse latent-space representation of actives, with highly potent larvicides confined to small, discontinuous regions, consistent with their broader entropy profile. As the activity threshold became less stringent, actives progressively expanded across multiple embedding neighborhoods and approached inactive regions, generating wider transition zones between classes at the 1 and 10 µM tasks. This organization helps explain the greater decision-boundary uncertainty observed for these intermediate thresholds. In the 100 µM task, the active class occupied a broader, more continuous latent-space domain, suggesting that the less stringent cutoff yielded a better-resolved active region.

Subsequently, the impact of predictive uncertainty on atom-level explanations derived from isosteric counterfactual perturbations was examined for the 10 µM task (Figure 3d). Four representative cases were prioritized for this analysis, comprising two compounds with higher uncertainty and two compounds with lower uncertainty. The higher-uncertainty cases were active compounds with intermediate predicted probabilities of 0.65 ± 0.22 and 0.68 ± 0.24, mutual information values of 0.11 and 0.14, and confidence margins of 0.64 and 0.36, respectively. These profiles indicate substantial stochastic variability in the predicted probability, which reduced the stability of the counterfactual attributions and produced less resolved substructural contribution patterns. By contrast, the lower-uncertainty cases, comprising one active and one inactive prediction, showed probabilities of 0.97 ± 0.01 and 0.00 ± 0.00, mutual information values ∼0.01, and confidence margins of 0.95 and 0.99, respectively. These stable predictions produced more coherent counterfactual maps, supporting a stronger association between the isosterically perturbed molecular regions and the predicted probability shifts. Therefore, high predictive uncertainty weakened the interpretability of atom-level counterfactual maps, whereas low-uncertainty predictions supported more reliable structure–activity relationships.

### 3.4. Virtual screening

A sequential virtual screening workflow was designed to prioritize novel larvicidal candidates from a commercially available library of 1.3 million compounds by applying our model as a threshold-dependent filter for progressive potency enrichment (Figure 4). Initially, the model trained on the 100 µM task was used as a permissive primary filter, reducing the library to approximately 100,000 compounds predicted to have larvicidal potential. Subsequently, the 10 µM task was applied as an intermediate potency filter, narrowing the set to 10,000 compounds. The more stringent 1 µM task was then used to further enrich the screening output toward highly potent candidates, yielding 500 compounds for downstream analysis. From these, chemical similarity analysis was performed both to exclude compounds previously tested in the literature and to enforce structural diversity, finally producing 10 virtual hits (see Table 1), representing a small and chemically diverse subset of candidates for experimental validation.

**Table 1.**
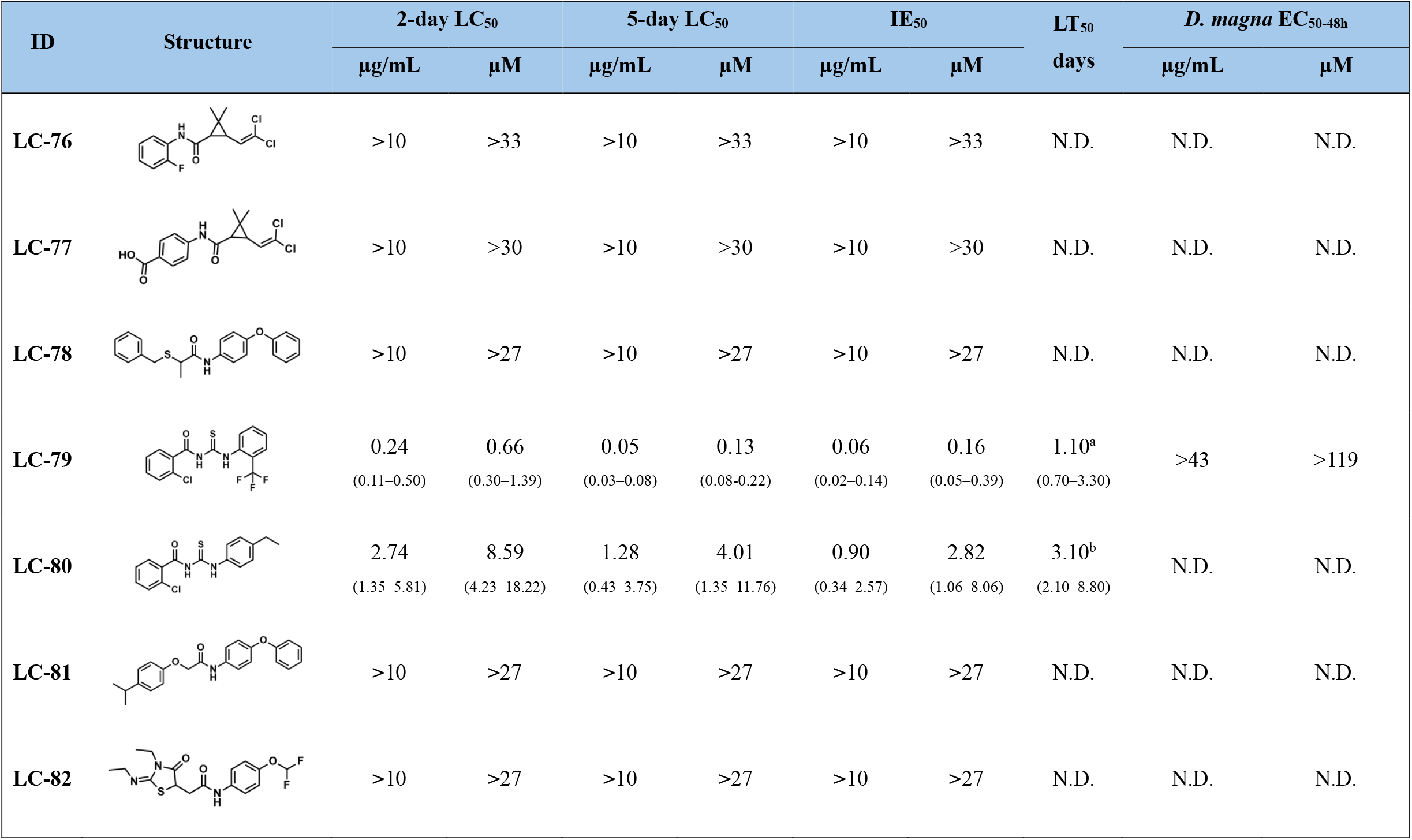
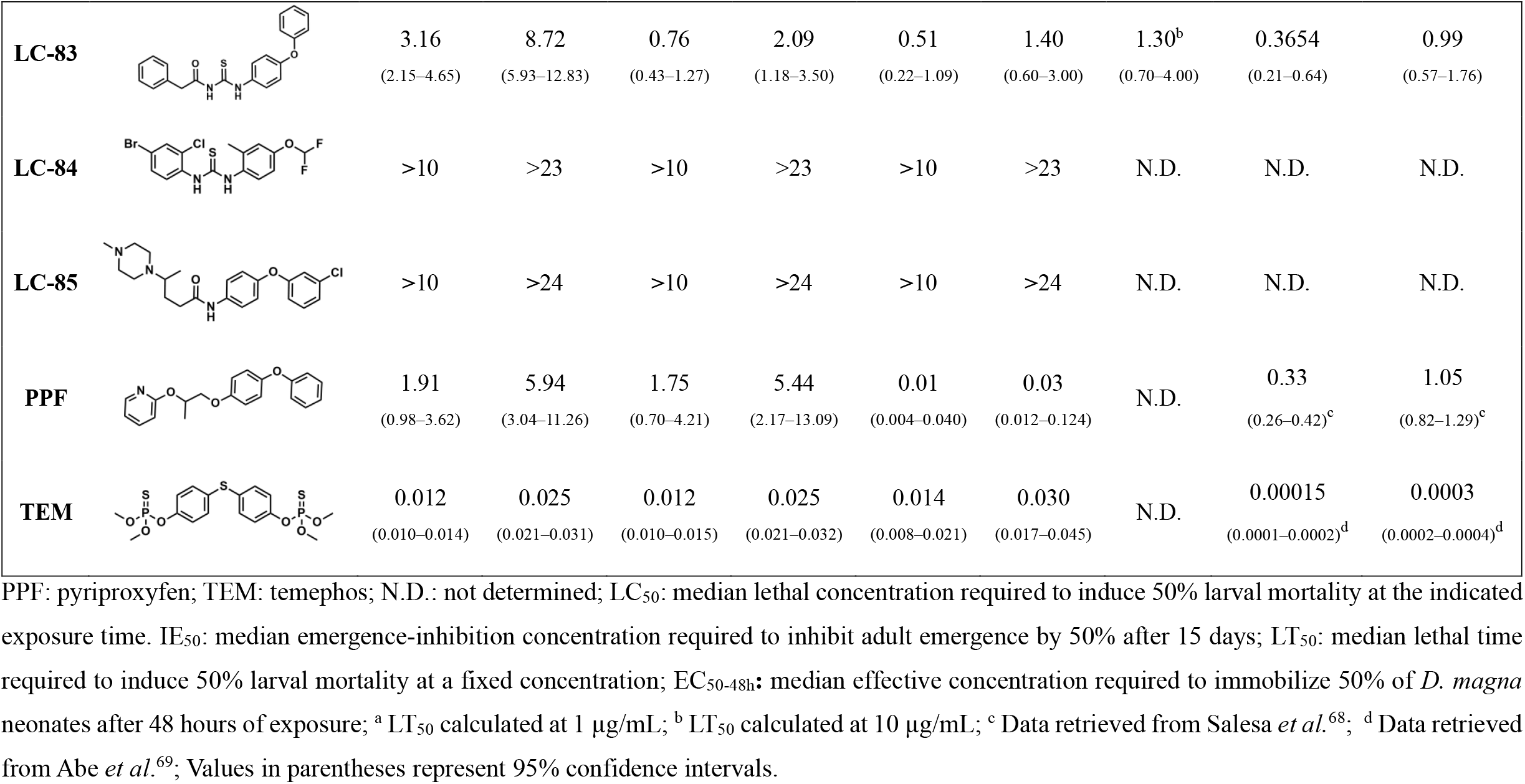
Chemical structures, larvicidal activity (2-day and 5-day LC_50_), developmental inhibition (IE_50_), and lethal time (LT_50_) of prioritized compounds against *Aedes aegypti* larvae, and their acute toxicity to *Daphnia magna*.

| ID | Structure | 2-day LC<sub>50</sub> |  | 5-day LC<sub>50</sub> |  | IE<sub>50</sub> |  | LT<sub>50</sub><br>days | *D. magna* EC<sub>50-48h</sub> |  |
| --- | --- | --- | --- | --- | --- | --- | --- | --- | --- | --- |
|  |  | µg/mL | µM | µg/mL | µM | µg/mL | µM |  | µg/mL | µM |
| LC-76 |  | >10 | >33 | >10 | >33 | >10 | >33 | N.D. | N.D. | N.D. |
| LC-77 |  | >10 | >30 | >10 | >30 | >10 | >30 | N.D. | N.D. | N.D. |
| LC-78 |  | >10 | >27 | >10 | >27 | >10 | >27 | N.D. | N.D. | N.D. |
| LC-79 |  | 0.24<br>(0.11–0.50) | 0.66<br>(0.30–1.39) | 0.05<br>(0.03–0.08) | 0.13<br>(0.08–0.22) | 0.06<br>(0.02–0.14) | 0.16<br>(0.05–0.39) | 1.10<sup>a</sup><br>(0.70–3.30) | >43 | >119 |
| LC-80 |  | 2.74<br>(1.35–5.81) | 8.59<br>(4.23–18.22) | 1.28<br>(0.43–3.75) | 4.01<br>(1.35–11.76) | 0.90<br>(0.34–2.57) | 2.82<br>(1.06–8.06) | 3.10<sup>b</sup><br>(2.10–8.80) | N.D. | N.D. |
| LC-81 |  | >10 | >27 | >10 | >27 | >10 | >27 | N.D. | N.D. | N.D. |
| LC-82 |  | >10 | >27 | >10 | >27 | >10 | >27 | N.D. | N.D. | N.D. |
| **LC-83** |  | 3.16<br>(2.15–4.65) | 8.72<br>(5.93–12.83) | 0.76<br>(0.43–1.27) | 2.09<br>(1.18–3.50) | 0.51<br>(0.22–1.09) | 1.40<br>(0.60–3.00) | 1.30<sup>b</sup><br>(0.70–4.00) | 0.3654<br>(0.21–0.64) | 0.99<br>(0.57–1.76) |
| **LC-84** |  | >10 | >23 | >10 | >23 | >10 | >23 | N.D. | N.D. | N.D. |
| **LC-85** |  | >10 | >24 | >10 | >24 | >10 | >24 | N.D. | N.D. | N.D. |
| **PPF** |  | 1.91<br>(0.98–3.62) | 5.94<br>(3.04–11.26) | 1.75<br>(0.70–4.21) | 5.44<br>(2.17–13.09) | 0.01<br>(0.004–0.040) | 0.03<br>(0.012–0.124) | N.D. | 0.33<br>(0.26–0.42)<sup>c</sup> | 1.05<br>(0.82–1.29)<sup>c</sup> |
| **TEM** |  | 0.012<br>(0.010–0.014) | 0.025<br>(0.021–0.031) | 0.012<br>(0.010–0.015) | 0.025<br>(0.021–0.032) | 0.014<br>(0.008–0.021) | 0.030<br>(0.017–0.045) | N.D. | 0.00015<br>(0.0001–0.0002)<sup>d</sup> | 0.0003<br>(0.0002–0.0004)<sup>d</sup> |

PPF: pyriproxyfen; TEM: temephos; N.D.: not determined; LC<sub>50</sub>: median lethal concentration required to induce 50% larval mortality at the indicated exposure time. IE<sub>50</sub>: median emergence-inhibition concentration required to inhibit adult emergence by 50% after 15 days; LT<sub>50</sub>: median lethal time required to induce 50% larval mortality at a fixed concentration; EC<sub>50-48h</sub>: median effective concentration required to immobilize 50% of *D. magna* neonates after 48 hours of exposure; <sup>a</sup> LT<sub>50</sub> calculated at 1 µg/mL; <sup>b</sup> LT<sub>50</sub> calculated at 10 µg/mL; <sup>c</sup> Data retrieved from Salesa *et al.*<sup>68</sup>; <sup>d</sup> Data retrieved from Abe *et al.*<sup>69</sup>; Values in parentheses represent 95% confidence intervals.

**Figure 4.**
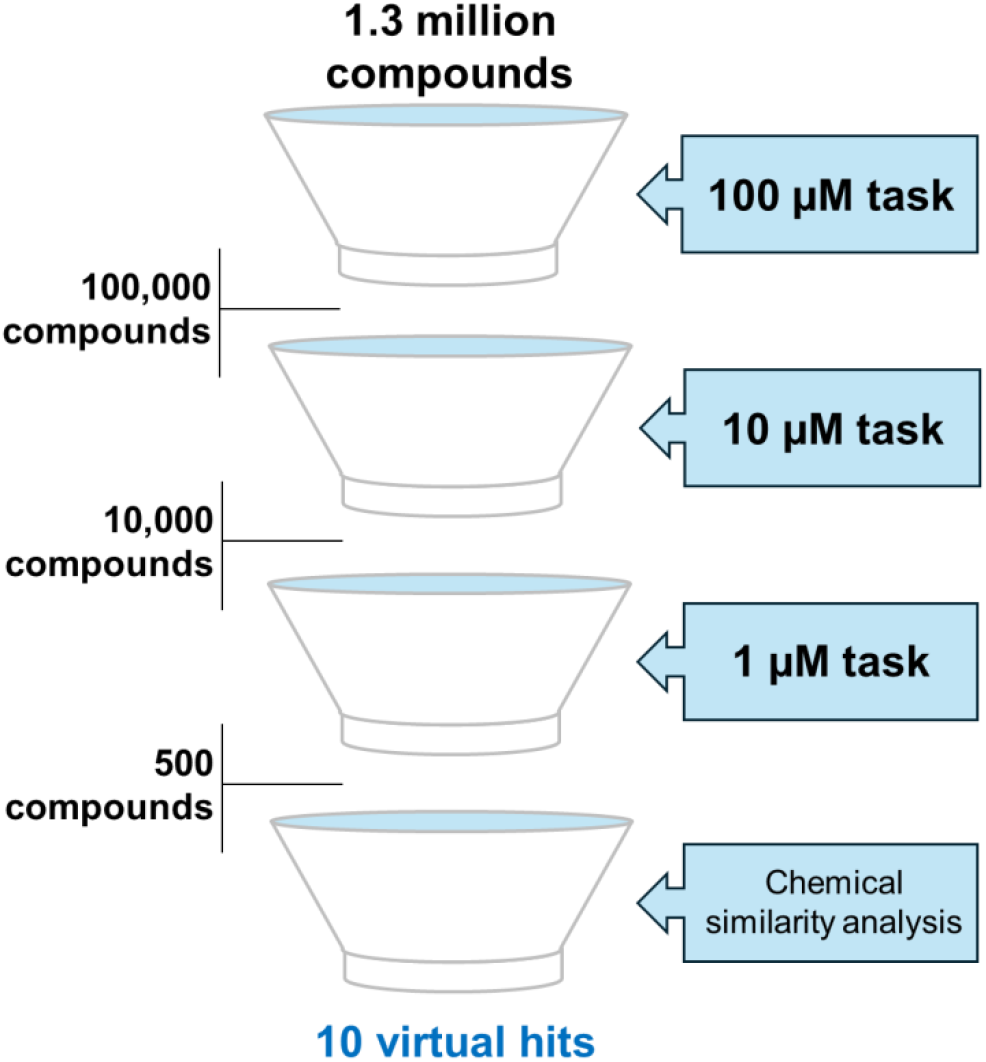
Virtual screening workflow for identifying novel larvicidal compounds.

This prioritization strategy produced a compact experimental test set with a measurable-hit rate of 30% under the tested conditions. Importantly, all three active compounds shared an acylthiourea-containing scaffold, suggesting that the workflow did not merely retrieve isolated actives but enriched a chemically coherent larvicidal chemotype. This result supports the practical utility of the sequential threshold-based screening strategy for reducing large commercial libraries to small, experimentally tractable candidate sets.

### 3.5 Larvicidal assays

The 10 putative hits were purchased and experimentally evaluated against third-instar *A. aegypti* larvae (Table 1). Among the selected compounds, three acylthiourea derivatives showed measurable larvicidal activity: 2-chloro-*N*-({[2-(trifluoromethyl)phenyl]amino}carbonothioyl)benzamide (**LC-79**), 2-chloro-*N*-{[(4-ethylphenyl)amino]carbonothioyl}benzamide (**LC-80**), and *N*-{[(4-phenoxyphenyl)amino]carbonothioyl}-2-phenylacetamide (**LC-83**). **LC-79** was the most potent hit, with 2-day and 5-day LC_50_ values of 0.24 µg/mL (0.66 µM) and 0.05 µg/mL (0.13 µM), respectively. **LC-80** and **LC-83** showed lower but consistent larvicidal activity, with 2-day LC_50_ values of 2.74 µg/mL (8.59 µM) and 3.16 µg/mL (8.72 µM), respectively. After 5 days, both compounds showed increased activity, with LC_50_ values of 1.28 µg/mL (4.01 µM) for LC-80 and 0.76 µg/mL (2.09 µM) for LC-83 (Table 1).

To further characterize the temporal dynamics of larval mortality, time-to-effect analysis was performed using the LT_50_ (see Table 1). Notably, **LC-79** showed the fastest larvicidal effect, with an LT_50_ of 1.10 days at 1 µg/mL. **LC-80** and **LC-83** also produced time-dependent mortality, with LT_50_ values of 3.10 and 1.30 days at 10 µg/mL, respectively. Subsequently, developmental inhibition was assessed by monitoring adult emergence over 15 days. **LC-79, LC-80**, and **LC-83** inhibited adult emergence with IE_50_ values of 0.06 µg/mL (0.16 µM), 0.90 µg/mL (2.82 µM), and 0.51 µg/mL (1.40 µM), respectively. These endpoint-level results positioned **LC-79** as the most competitive validated hit. **LC-79** reached the same submicromolar order of magnitude as temephos after 5 days and outperformed pyriproxyfen in larval mortality, with approximately 9-fold and 42-fold greater molar potency at 2 and 5 days, respectively.

Structurally, the shared acylthiourea motif observed among the active hits provides a sulfur-substituted counterpart to the urea linkage found in acylurea insect growth regulators. This structural relationship provides a rationale for benchmarking **LC-79** against urea-related larvicidal scaffolds. Compared with the acylthiourea-containing anthranilic diamides reported by Kavyasri *et al*.,^70^ **LC-79** showed a stronger larvicidal profile under the same exposure window. **LC-79** also reached the same activity range reported for novaluron, a benzoylurea insect growth regulator with a 7-day LC_50_ of 0.047 µg/mL against *A. aegypti*.^71^ Chemically, the C=O-to-C=S replacement may reduce hydrogen-bond acceptor strength while increasing sulfur polarizability, thereby modulating lipophilic balance and desolvation relative to acylurea scaffolds. This feature offers a useful physicochemical handle for further optimization of the **LC-79** scaffold and supports its development as a chemically distinct larvicidal hit.

The activity differences among **LC-79, LC-80**, and **LC-83** also provide an initial SAR hypothesis for this chemotype. **LC-79** differs from the less potent analogs by combining the acylthiourea linker with a chloro-substituted benzamide region and a trifluoromethyl-substituted anilide moiety. This substitution pattern may improve larval penetration, binding-site complementarity, or metabolic stability relative to the ethylphenyl and phenoxyphenyl analogs. Although additional analogs are required to confirm these trends, the current hit series suggests that halogenated and electron-withdrawing aryl substituents may be favorable optimization handles for increasing acylthiourea larvicidal potency. Nevertheless, these structural comparisons should not be interpreted as evidence that **LC-79** acts through the same molecular mechanism as benzoylurea or acylurea insect growth regulators. The observed larval mortality, rapid time-to-effect, and adult-emergence inhibition indicate that **LC-79** affects both survival and development, but target-level mechanism-of-action studies were not performed in the present work. Future studies should therefore evaluate whether **LC-79** interferes with chitin biosynthesis, endocrine regulation, mitochondrial function, neuromuscular signaling, or other larval physiological processes.

### 3.6. Acute toxicity assay

Acute toxicity to *D. magna* was subsequently assessed to estimate the selectivity of the validated hits over a representative non-target aquatic organism. *D. magna* was included as a standardized and ecologically relevant non-target aquatic model, since larvicidal compounds intended for mosquito control may ultimately reach freshwater environments and affect aquatic invertebrates. As shown in Table 1, **LC-79** showed no measurable acute toxicity to *D. magna* at the highest tested concentration, with an EC_50-48h_ >43 µg/mL (>119 µM), corresponding to selectivity indices >180 and >860 relative to its 2-day and 5-day LC_50_ values, respectively. This selectivity window was markedly broader than those reported for the positive controls. Pyriproxyfen showed a reported *D. magna* EC_50-48h_ of 0.33 µg/mL (1.05 µM),^68^ corresponding to selectivity indices of 0.18 and 0.19 relative to its 2-day and 5-day LC_50_ values, respectively. Temephos showed substantially higher acute toxicity to *D. magna*,^69^ with a reported EC_50-48h_ of 0.00015 µg/mL (0.0003 µM) and selectivity indices of approximately 0.01 for both larvicidal endpoints. By contrast, **LC-83** showed an EC_50_ of 0.36 µg/mL (0.99 µM) against *D. magna*, corresponding to selectivity indices of 0.11 and 0.47 relative to its 2-day and 5-day LC_50_ values, respectively. Although **LC-83** and pyriproxyfen displayed similar acute toxicity ranges in *D. magna*, only **LC-79** combined submicromolar larvicidal activity with a wide preliminary selectivity margin over this non-target aquatic organism. These results reinforce **LC-79** as the most promising validated hit, combining submicromolar LC_50_ values, rapid larval mortality, potent inhibition of adult emergence at submicromolar concentrations, and low acute toxicity to *D. magna* under the tested conditions.

These findings should be interpreted as an encouraging preliminary selectivity signal rather than a complete environmental safety profile. *D. magna* provides a relevant and widely used aquatic non-target model, but additional assays will be required to define the ecotoxicological window of **LC-79**, including chronic exposure, effects on other aquatic invertebrates and fish, degradation behavior, photostability, sediment partitioning, and activity under environmentally realistic water conditions.

Several limitations should be considered when interpreting these findings. First, the predictive models were trained on a relatively small curated dataset assembled from heterogeneous public sources, and LC_50_-derived threshold labels necessarily simplify continuous potency information.

Second, although scaffold splitting provided a stricter estimate of chemical-space generalization, prospective performance may vary for chemotypes outside the training-domain distribution. Third, the experimental validation was performed under laboratory conditions using a single *A. aegypti* colony; therefore, activity against geographically diverse and insecticide-resistant populations remains to be determined. Fourth, **LC-79** showed a favorable acute *D. magna* selectivity profile, but broader ecotoxicological evaluation is required before environmental safety can be inferred. Finally, the molecular mechanism responsible for **LC-79**-mediated larval mortality and adult-emergence inhibition remains unknown. Addressing these limitations through resistance-strain testing, semi-field assays, target-deconvolution studies, and expanded non-target toxicity profiling will be essential for advancing **LC-79** from validated hit to optimized larvicide lead.

## 4. Conclusion

In this study, we developed and experimentally validated an instance-wise contrastive GNN workflow for larvicidal hit discovery against Aedes aegypti. The model achieved strong predictive performance, outperformed conventional machine-learning and graph-based baselines, and retained useful performance under scaffold-based evaluation, supporting its application as a prospective enrichment tool. Importantly, the workflow moved beyond retrospective modeling by screening 1.3 million commercially available compounds and prioritizing 10 candidates for biological testing, of which three showed measurable larvicidal activity. Among these, **LC-79** emerged as the most promising candidate, combining potent larvicidal activity against *A. aegypti* larvae, with 2-day and 5-day LC_50_ values of 0.24 µg/mL (0.66 µM) and 0.05 µg/mL (0.13 µM), respectively, rapid time-to-effect (LT_50_ = 1.10 days at 1 µg/mL), and submicromolar inhibition of adult emergence [IE_50_ = 0.06 µg/mL (0.16 µM)]. Importantly, **LC-79** showed no measurable acute toxicity to *D. magna* at the highest tested concentration [EC_50-48h_ >43 µg/mL (>119 µM)], resulting in selectivity indices >180 and >915 relative to its 2-day and 5-day larvicidal endpoints, respectively. Structurally, **LC-79** introduces an acylthiourea scaffold distinct from classical benzoylurea insect growth regulators, highlighting a new chemotype for larvicide discovery. Overall, these findings support **LC-79** as a validated larvicidal hit for prospective semi-field and field studies, as well as more detailed ecotoxicological and structure–activity optimization campaigns.

## Data and Software Availability

The curated datasets and source code generated for this study are publicly available in the GitHub repository: https://github.com/TaoLen/AedesGNN.

## Author Contributions

All authors contributed to the study’s conception and design. K.S.L.C., R.C.B., E.M., G.A.R.O., and B.J.N. conceived and designed the study. K.S.L.C. performed data collection, curation, and dataset preparation. K.S.L.C., V.A.F.C., A.S.S., R.C.B., and B.J.N. performed computational modeling, software implementation, and virtual screening. G.H.G.C., C.L., and J.R. performed the larvicidal assays. A.S.S., C.S.P., B.C.B., L.B.M., and G.A.R.O. performed the acute toxicity assays. K.S.L.C., G.H.G.C., V.A.F.C., A.S.S., G.A.R.O., C.L., J.R., and B.J.N. analyzed and interpreted the data. V.A.F.C., A.S.S., G.A.R.O., J.R., and B.J.N. wrote the original draft. J.R., H.M., R.C.B., E.M., S.S., G.A.R.O., and C.L. reviewed and edited the manuscript. G.A.R.O., C.L., J.R., and B.J.N. supervised the study. S.S. and B.J.N. secured funding and managed the project. All authors approved the final manuscript and ensured the study’s accuracy and integrity.

## Notes

Rodolpho de Campos Braga and Eugene N. Muratov are co-founders of InsilicAll Ltda. and Predictive LLC, respectively, which develop novel alternative methods and software for toxicity prediction. All the other authors declare no conflicts.

## Acknowledgements

The authors thank the Conselho Nacional de Desenvolvimento Científico e Tecnológico (CNPq), the Coordenação de Aperfeiçoamento de Pessoal de Nível Superior (CAPES), and the Fundação de Amparo à Pesquisa do Estado de Goiás (FAPEG) for financial support and fellowships. The authors also acknowledge Emival Sebastião de Carvalho for his institutional support and efforts that contributed to the establishment of the Laboratory of Cheminformatics.

## Supporting Information

The Supporting Information contains detailed statistical performance results for instance-wise contrastive GNN under random and scaffold split strategies, as well as for baseline models comprising SVM, Random Forest, MPNN, GAT, GIN, and AttentiveFP.

## References

(1) World Health Organization. Vector-borne diseases. https://www.who.int/news-room/fact-sheets/detail/vector-borne-diseases (xaccessed 2026-05-06).

(2) Paz-Bailey, G.; Adams, L. E.; Deen, J.; Anderson, K. B.; Katzelnick, L. C. Dengue. The Lancet 2024, 403 (10427), 667–682. 10.1016/S0140-6736(23)02576-X.

(3) de Souza, W. M.; Lecuit, M.; Weaver, S. C. Chikungunya Virus and Other Emerging Arthritogenic Alphaviruses. Nat. Rev. Microbiol. 2025, 23 (9), 585–601. 10.1038/s41579-025-01177-8.

(4) Brasil, P.; Nielsen-Saines, K.; Guaraldo, L.; Fuller, T.; Moreira, M. E. L. A Decade Later, What Have We Learned from the Zika Epidemic in Children with Intrauterine Exposure? The Lancet 2025, 406 (10500), 295–306. 10.1016/S0140-6736(25)00826-8.

(5) Ortiz-Prado, E.; Prieto-Marin, J. G.; Izquierdo-Condoy, J. S.; Vasconez-Gonzalez, J.; Villamil-Parra, W. A.; Viscor, G.; Niño-Méndez, Ó. A.; Correa-Bautista, J. E.; Rusiñol, M.; Cevallos-Robalino, D.; Navarro, J.-C.; Villalobos-Madriz, J. A. Yellow Fever in Latin America and the Escalating Risks in a Changing Eco-Epidemiological Landscape: A Review. Lancet Reg. Health Am. 2026, 56, 101431. 10.1016/j.lana.2026.101431.

(6) World Health Organization. Dengue: global situation, surveillance and progress – 2024 update. https://www.who.int/publications/i/item/who-wer10052-665-678 (accessed 2026-05-06).

(7) Xu, Z.; Bambrick, H.; Frentiu, F. D.; Devine, G.; Yakob, L.; Williams, G.; Hu, W. Projecting the Future of Dengue under Climate Change Scenarios: Progress, Uncertainties and Research Needs. PLoS Negl. Trop. Dis. 2020, 14 (3), e0008118. 10.1371/journal.pntd.0008118.

(8) Colón-González, F. J.; Sewe, M. O.; Tompkins, A. M.; Sjödin, H.; Casallas, A.; Rocklöv, J.; Caminade, C.; Lowe, R. Projecting the Risk of Mosquito-Borne Diseases in a Warmer and More Populated World: A Multi-Model, Multi-Scenario Intercomparison Modelling Study. Lancet Planet. Health 2021, 5 (7), e404–e414. 10.1016/S2542-5196(21)00132-7.

(9) Barcellos, C.; Matos, V.; Lana, R. M.; Lowe, R. Climate Change, Thermal Anomalies, and the Recent Progression of Dengue in Brazil. Sci. Rep. 2024, 14 (1), 5948. 10.1038/s41598-024-56044-y.

(10) Roiz, D.; Wilson, A. L.; Scott, T. W.; Fonseca, D. M.; Jourdain, F.; Müller, P.;Velayudhan, R.; Corbel, V. Integrated Aedes Management for the Control of Aedes-Borne Diseases. PLoS Negl. Trop. Dis. 2018, 12 (12), e0006845. 10.1371/journal.pntd.0006845.

(11) Oxborough, R. M.; Emidi, B.; Yougang, A. P.; Abeku, T. A.; Ahmed, F.; Biggs, J. R.; Chan, K.; Cook, J.; Edwards, A.; Falconer, J.; Kamgang, B.; Messenger, L. A.; Seelig, F.; Taylor, R.; Tedjou, A. N.; Lines, J.; Clarke, S. E.; Kristan, M. Building Resilience against the Growing Threat of Arboviruses: A Scoping Review of Aedes Vector Surveillance, Control Strategies and Insecticide Resistance in Africa. Parasit. Vectors 2025, 18 (1), 415. 10.1186/s13071-025-07049-7.

(12) Marcombe, S.; Chonephetsarath, S.; Thammavong, P.; Brey, P. T. Alternative Insecticides for Larval Control of the Dengue Vector Aedes Aegypti in Lao PDR: Insecticide Resistance and Semi-Field Trial Study. Parasit. Vectors 2018, 11 (1), 616. 10.1186/s13071-018-3187-8.

(13) Marina, C. F.; Bond, J. G.; Muñoz, J.; Valle, J.; Quiroz-Martínez, H.; Torres-Monzón, J. A.; Williams, T.; Marina, C. F.; Bond, J. G.; Muñoz, J.; Valle, J.; Quiroz-Martínez, H.; Torres-Monzón, J. A.; Williams, T. Comparison of Novaluron, Pyriproxyfen, Spinosad and Temephos as Larvicides against Aedes Aegypti in Chiapas, Mexico. Salud Pública México 2020, 62 (4), 424–431. 10.21149/10168.

(14) Davila-Barboza, J. A.; Gutierrez-Rodriguez, S. M.; Juache-Villagrana, A. E.; Lopez-Monroy, B.; Flores, A. E. Widespread Resistance to Temephos in Aedes Aegypti (Diptera: Culicidae) from Mexico. Insects 2024, 15 (2), 120. 10.3390/insects15020120.

(15) Bellinato, D. F.; Viana-Medeiros, P. F.; Araújo, S. C.; Martins, A. J.; Lima, J. B. P.; Valle, D. Resistance Status to the Insecticides Temephos, Deltamethrin, and Diflubenzuron in Brazilian Aedes Aegypti Populations. BioMed Res. Int. 2016, 2016, 8603263. 10.1155/2016/8603263.

(16) Campos, K. B.; Alomar, A. A.; Eastmond, B. H.; Obara, M. T.; S Dias, L. D.; Rahman, R. U.; Alto, B. W. Assessment of Insecticide Resistance of Aedes Aegypti (Diptera: Culicidae) Populations to Insect Growth Regulator Pyriproxyfen, in the Northeast Region of Brazil. J. Vector Ecol. J. Soc. Vector Ecol. 2023, 48 (1), 12–18. 10.52707/1081-1710-48.1.12.

(17) Campos, K. B.; Alomar, A. A.; Eastmond, B. H.; Obara, M. T.; Alto, B. W. Brazilian Populations of Aedes Aegypti Resistant to Pyriproxyfen Exhibit Lower Susceptibility to Infection with Zika Virus. Viruses 2022, 14 (10), 2198.10.3390/v14102198.

(18) Montaño-Reyes, A.; Llanderal-Cázares, C.; Valdez-Carrasco, J.; Miranda-Perkins, K.; Sánchez-Arroyo, H. Susceptibility and Alterations by Diflubenzuron in Larvae of Aedes Aegypti. Arch. Insect Biochem. Physiol. 2019, 102 (2), e21604. 10.1002/arch.21604.

(19) Tominaga, F. K.; Brito, R. S.; Oliveira do Nascimento, J.; Giannocco, G.; Monteiro de Barros Maciel, R.; Kummrow, F.; Pereira, B. F. Pyriproxyfen Toxicity to Fish and Crustaceans: A Literature Review. Environ. Res. 2025, 274, 121295. 10.1016/j.envres.2025.121295.

(20) Vieira Santos, V. S.; Caixeta, E. S.; Campos Júnior, E. O. de; Pereira, B. B. Ecotoxicological Effects of Larvicide Used in the Control of Aedes Aegypti on Nontarget Organisms: Redefining the Use of Pyriproxyfen. J. Toxicol. Environ. Health A 2017, 80 (3), 155–160. 10.1080/15287394.2016.1266721.

(21) Truong, L.; Gonnerman, G.; Simonich, M. T.; Tanguay, R. L. Assessment of the Developmental and Neurotoxicity of the Mosquito Control Larvicide, Pyriproxyfen, Using Embryonic Zebrafish. Environ. Pollut. Barking Essex 1987 2016, 218, 1089–1093. 10.1016/j.envpol.2016.08.061.

(22) Teixeira, J. R. de S.; de Souza, A. M.; Macedo-Sampaio, J. V. de Menezes, F. P.; Pereira, B. F.; de Medeiros, S. R. B.; Luchiari, A. C. Embryotoxic Effects of Pesticides in Zebrafish (Danio Rerio): Diflubenzuron, Pyriproxyfen, and Its Mixtures. Toxics 2024, 12 (2), 160. 10.3390/toxics12020160.

(23) Benitez-Trinidad, A. B.; Herrera-Moreno, J. F.; Vázquez-Estrada, G.; Verdín-Betancourt, F. A.; Sordo, M.; Ostrosky-Wegman, P.; Bernal-Hernández, Y. Y.; Medina-Díaz, I. M.; Barrón-Vivanco, B. S.; Robledo-Marenco, M. L.; Salazar, A. M.; Rojas-García, E. Cytostatic and Genotoxic Effect of Temephos in Human Lymphocytes and HepG2 Cells. Toxicol. In Vitro 2015, 29 (4), 779–786. 10.1016/j.tiv.2015.02.008.

(24) Bao, Y.; Chen, Y.; Zhou, Y.; Wang, Q.; Zuo, Z.; Yang, C. Chronic Diflubenzuron Exposure Causes Reproductive Toxic Effects in Female Marine Medaka (Oryzias Melastigma). Aquat. Toxicol. 2023, 258, 106511. 10.1016/j.aquatox.2023.106511.

(25) Kong, Y.; Zhou, C.; Tan, D.; Xu, X.; Li, Z.; Cheng, J. Discovery of Potential Neonicotinoid Insecticides by an Artificial Intelligence Generative Model and Structure-Based Virtual Screening. J. Agric. Food Chem. 2024, 72 (10), 5145–5152. 10.1021/acs.jafc.3c06895.

(26) Djoumbou-Feunang, Y.; Wilmot, J.; Kinney, J.; Chanda, P.; Yu, P.; Sader, A.; Sharifi, M.; Smith, S.; Ou, J.; Hu, J.; Shipp, E.; Tomandl, D.; Kumpatla, S. P. Cheminformatics and Artificial Intelligence for Accelerating Agrochemical Discovery. Front. Chem. 2023, 11, 1292027. 10.3389/fchem.2023.1292027.

(27) Sparks, T. C. Trends in Insecticide Discovery: A Review, Analysis and Perspective. Pestic. Biochem. Physiol. 2025, 213, 106521. 10.1016/j.pestbp.2025.106521.

(28) Cherkasov, A.; Muratov, E. N.; Fourches, D.; Varnek, A.; Baskin, I. I.; Cronin, M.; Dearden, J.; Gramatica, P.; Martin, Y. C.; Todeschini, R.; Consonni, V.; Kuz’min, V. E.; Cramer, R.; Benigni, R.; Yang, C.; Rathman, J.; Terfloth, L.; Gasteiger, J.; Richard, A.; Tropsha, A. QSAR Modeling: Where Have You Been? Where Are You Going To? J. Med. Chem. 2014, 57 (12), 4977–5010. 10.1021/jm4004285.

(29) Soares, T. A.; Nunes-Alves, A.; Mazzolari, A.; Ruggiu, F.; Wei, G.-W.; Merz, K. The (Re)-Evolution of Quantitative Structure–Activity Relationship (QSAR) Studies Propelled by the Surge of Machine Learning Methods. J. Chem. Inf. Model. 2022, 62 (22), 5317–5320. 10.1021/acs.jcim.2c01422.

(30) Saavedra, L. M.; Romanelli, G. P.; Duchowicz, P. R. Quantitative Structure-Activity Relationship (QSAR) Analysis of Plant-Derived Compounds with Larvicidal Activity against Zika Aedes Aegypti (Diptera: Culicidae) Vector Using Freely Available Descriptors. Pest Manag. Sci. 2018, 74 (7), 1608–1615. 10.1002/ps.4850.

(31) Saavedra, L. M.; Romanelli, G. P.; Rozo, C. E.; Duchowicz, P. R. The Quantitative Structure–Insecticidal Activity Relationships from Plant Derived Compounds against Chikungunya and Zika Aedes Aegypti (Diptera:Culicidae) Vector. Sci. Total Environ. 2018, 610–611, 937–943. 10.1016/j.scitotenv.2017.08.119.

(32) Saavedra, L. M.; Romanelli, G. P.; Duchowicz, P. R. A Non-Conformational QSAR Study for Plant-Derived Larvicides against Zika Aedes Aegypti L. Vector. Environ. Sci. Pollut. Res. 2020, 27 (6), 6205–6214. 10.1007/s11356-019-06630-9.

(33) Gomes, J. P. A.; Mourão, E. D. S.; dos Anjos, J. V.; de Alencar Filho, E. B. QSAR Modelling and Structural Aspects Concerning Synthetic Heterocycles with Larvicidal Activity against Aedes Aegypti. Struct. Chem. 2020, 31 (6), 2501–2512.10.1007/s11224-020-01597-7.

(34) Devillers, J.; Doucet-Panaye, A.; Doucet, J. P. Structure-Activity Relationship (SAR) Modelling of Mosquito Larvicides. SAR QSAR Environ. Res. 2015, 26 (4), 263–278. 10.1080/1062936X.2015.1026571.

(35) Wigh, D. S.; Goodman, J. M.; Lapkin, A. A. A Review of Molecular Representation in the Age of Machine Learning. WIREs Comput. Mol. Sci. 2022, 12 (5), e1603. 10.1002/wcms.1603.

(36) Chuang, K. V.; Gunsalus, L. M.; Keiser, M. J. Learning Molecular Representations for Medicinal Chemistry. J. Med. Chem. 2020, 63 (16), 8705–8722. 10.1021/acs.jmedchem.0c00385.

(37) Reiser, P.; Neubert, M.; Eberhard, A.; Torresi, L.; Zhou, C.; Shao, C.; Metni, H.; van Hoesel, C.; Schopmans, H.; Sommer, T.; Friederich, P. Graph Neural Networks for Materials Science and Chemistry. Commun. Mater. 2022, 3 (1), 93. 10.1038/s43246-022-00315-6.

(38) Zhang, O.; Lin, H.; Zhang, X.; Wang, X.; Wu, Z.; Ye, Q.; Zhao, W.; Wang, J.; Ying, K.; Kang, Y.; Hsieh, C.-Y.; Hou, T. Graph Neural Networks in Modern AI-Aided Drug Discovery. Chem. Rev. 2025, 125 (20), 10001–10103. 10.1021/acs.chemrev.5c00461.

(39) Wieder, O.; Kohlbacher, S.; Kuenemann, M.; Garon, A.; Ducrot, P.; Seidel, T.; Langer, T. A Compact Review of Molecular Property Prediction with Graph Neural Networks. Drug Discov. Today Technol. 2020, 37, 1–12. 10.1016/j.ddtec.2020.11.009.

(40) Sultan, A.; Sieg, J.; Mathea, M.; Volkamer, A. Transformers for Molecular Property Prediction: Lessons Learned from the Past Five Years. J. Chem. Inf. Model. 2024, 64 (16), 6259–6280. 10.1021/acs.jcim.4c00747.

(41) Santos, E. S. de A.; Sandes, G. F. S.; da Silva, A. C. G.; Martin, H.-J.; Muratov, E. N.; Braga, R. de C.; Neves, B. J. Multimodal Cross-Attentive Graph-Based Framework for Predicting In Vivo Endocrine Disruptors. Adv. Sci. 2026, 13 (21), e19897. 10.1002/advs.202519897.

(42) Gaulton, A.; Bellis, L. J.; Bento, A. P.; Chambers, J.; Davies, M.; Hersey, A.; Light, Y.; McGlinchey, S.; Michalovich, D.; Al-Lazikani, B.; Overington, J. P. ChEMBL: A Large-Scale Bioactivity Database for Drug Discovery. Nucleic Acids Res. 2012, 40 (Database issue), D1100–D1107. 10.1093/nar/gkr777.

(43) Mendez, D.; Gaulton, A.; Bento, A. P.; Chambers, J.; De Veij, M.; Félix, E.; Magariños, M. P.; Mosquera, J. F.; Mutowo, P.; Nowotka, M.; Gordillo-Marañón, M.; Hunter, F.; Junco, L.; Mugumbate, G.; Rodriguez-Lopez, M.; Atkinson, F.; Bosc, N.; Radoux, C. J.; Segura-Cabrera, A.; Hersey, A.; Leach, A. R. ChEMBL: Towards Direct Deposition of Bioassay Data. Nucleic Acids Res. 2019, 47 (D1), D930–D940. 10.1093/nar/gky1075.

(44) Olker, J. H.; Elonen, C. M.; Pilli, A.; Anderson, A.; Kinziger, B.; Erickson, S.; Skopinski, M.; Pomplun, A.; LaLone, C. A.; Russom, C. L.; Hoff, D. The ECOTOXicology Knowledgebase: A Curated Database of Ecologically Relevant Toxicity Tests to Support Environmental Research and Risk Assessment. Environ. Toxicol. Chem. 2022, 41 (6), 1520–1539. 10.1002/etc.5324.

(45) Fourches, D.; Muratov, E.; Tropsha, A. Trust, but Verify: On the Importance of Chemical Structure Curation in Cheminformatics and QSAR Modeling Research. J. Chem. Inf. Model. 2010, 50 (7), 1189–1204. 10.1021/ci100176x.

(46) Fourches, D.; Muratov, E.; Tropsha, A. Trust, but Verify II: A Practical Guide to Chemogenomics Data Curation. J. Chem. Inf. Model. 2016, 56 (7), 1243–1252. 10.1021/acs.jcim.6b00129.

(47) Sander, T.; Freyss, J.; von Korff, M.; Rufener, C. DataWarrior: An Open-Source Program For Chemistry Aware Data Visualization And Analysis. J. Chem. Inf. Model. 2015, 55 (2), 460–473. 10.1021/ci500588j.

(48) Wu, C.; Wu, F.; Qi, T.; Huang, Y.; Xie, X. Fastformer: Additive Attention Can Be All You Need. August 20, 2021. 10.48550/arXiv.2108.09084.

(49) Vaswani, A.; Shazeer, N.; Parmar, N.; Uszkoreit, J.; Jones, L.; Gomez, A. N.; Kaiser, Ł. ukasz; Polosukhin, I. Attention Is All You Need. In Advances in Neural Information Processing Systems; Curran Associates, Inc., 2017; Vol. 30, pp 1–11.

(50) Wang, Y.; Magar, R.; Liang, C.; Barati Farimani, A. Improving Molecular Contrastive Learning via Faulty Negative Mitigation and Decomposed Fragment Contrast. J. Chem. Inf. Model. 2022, 62 (11), 2713–2725. 10.1021/acs.jcim.2c00495.

(51) Chen, T.; Kornblith, S.; Norouzi, M.; Hinton, G. A Simple Framework for Contrastive Learning of Visual Representations. arXiv.org. https://arxiv.org/abs/2002.05709v3 (accessed 2026-05-07).

(52) Rogers, D.; Hahn, M. Extended-Connectivity Fingerprints. J. Chem. Inf. Model. 2010, 50 (5), 742–754. 10.1021/ci100050t.

(53) Degen, J.; Wegscheid-Gerlach, C.; Zaliani, A.; Rarey, M. On the Art of Compiling and Using “Drug-Like” Chemical Fragment Spaces. ChemMedChem 2008, 3 (10), 1503–1507. 10.1002/cmdc.200800178.

(54) Cipolla, R.; Gal, Y.; Kendall, A. Multi-Task Learning Using Uncertainty to Weigh Losses for Scene Geometry and Semantics. In 2018 IEEE/CVF Conference on Computer Vision and Pattern Recognition; 2018; pp 7482–7491. 10.1109/CVPR.2018.00781.

(55) Gal, Y.; Ghahramani, Z. Dropout as a Bayesian Approximation: Representing Model Uncertainty in Deep Learning. arXiv October 4, 2016. 10.48550/arXiv.1506.02142.

(56) Kendall, A.; Gal, Y. What Uncertainties Do We Need in Bayesian Deep Learning for Computer Vision? arXiv October 5, 2017. 10.48550/arXiv.1703.04977.

(57) Smith, L.; Gal, Y. Understanding Measures of Uncertainty for Adversarial Example Detection. arXiv March 22, 2018. https://arxiv.org/abs/1803.08533v1.

(58) Veličković, P.; Cucurull, G.; Casanova, A.; Romero, A.; Liò, P.; Bengio, Y. Graph Attention Networks. arXiv February 4, 2018. 10.48550/arXiv.1710.10903.

(59) Xiong, Z.; Wang, D.; Liu, X.; Zhong, F.; Wan, X.; Li, X.; Li, Z.; Luo, X.; Chen, K.; Jiang, H.; Zheng, M. Pushing the Boundaries of Molecular Representation for Drug Discovery with the Graph Attention Mechanism. J. Med. Chem. 2020, 63 (16), 8749–8760. 10.1021/acs.jmedchem.9b00959.

(60) Gilmer, J.; Schoenholz, S. S.; Riley, P. F.; Vinyals, O.; Dahl, G. E. Neural Message Passing for Quantum Chemistry. arXiv June 12, 2017. 10.48550/arXiv.1704.01212.

(61) Xu, K.; Hu, W.; Leskovec, J.; Jegelka, S. How Powerful Are Graph Neural Networks? arXiv February 22, 2019. 10.48550/arXiv.1810.00826.

(62) Costa, V. A. F.; Carvalho, A. M. dos S.; Chelucci, R. C.; Mendonça de Melo, F. da S.; Felizardo, G. S. S.; Bernardes, C. A. C.; Martin, H.-J.; Braga, R. de C.; Charneau, S.; Muratov, E. N.; Andricopulo, A. D.; Bastos, I. M. D.; Neves, B. J. Multiscale-Aware Graph Embedding Approach Uncovers LC-61, a Potent Anti-Leishmania Infantum Compound. J. Chem. Inf. Model. 2026, 66 (7), 3643–3659. 10.1021/acs.jcim.5c02947.

(63) Rodrigues, J.; Borges, P. R.; Fernandes, É. K. K.; Luz, C. Activity of Additives and Their Effect in Formulations of Metarhizium Anisopliae s.l. IP 46 against Aedes Aegypti Adults and on Post Mortem Conidiogenesis. Acta Trop. 2019, 193, 192–198. 10.1016/j.actatropica.2019.03.002.

(64) Lima, W. P.; Chiaravalloti Neto, F.; Macoris, M. de L. da G.; Zuccari, D. A. P. de C.; Dibo, M. R. Establishment of the feeding methodology of Aedes aegypti (Diptera-Culicidae) in Swiss mice and evaluation of the toxicity and residual effect of essential oil from Tagetes minuta L (Asteraceae), in populations of Aedes aegypti. Rev. Soc. Bras. Med. Trop. 2009, 42 (6), 638–641. 10.1590/s0037-86822009000600005.

(65) World Health Organization. Guidelines for Laboratory and Field Testing of Mosquito Larvicides, 2005. https://www.who.int/publications/i/item/WHO-CDS-WHOPES-GCDPP-2005.13 (accessed 2026-04-15).

(66) Organisation for Economic Co-operation and Development. Test No. 202: Daphnia Sp. Acute Immobilisation Test. OECD Guidel. Test. Chem. Sect. 2 2004. 10.1787/9789264069947-en.

(67) Vaswani, A.; Shazeer, N.; Parmar, N.; Uszkoreit, J.; Jones, L.; Gomez, A. N.; Kaiser, L.; Polosukhin, I. Attention Is All You Need. arXiv August 2, 2023. 10.48550/arXiv.1706.03762.

(68) Salesa, B.; Torres-Gavilá, J.; Sancho, E.; Ferrando, M. D. Multigenerational Effects of the Insecticide Pyriproxyfen and Recovery in Daphnia Magna. Sci. Total Environ. 2023, 886, 164013. 10.1016/j.scitotenv.2023.164013.

(69) Abe, F. R.; Coleone, A. C.; Machado, A. A.; Gonçalves Machado-Neto, J. Ecotoxicity and Environmental Risk Assessment of Larvicides Used in the Control of Aedes Aegypti to Daphnia Magna (Crustacea, Cladocera). J. Toxicol. Environ. Health A 2014, 77 (1–3), 37–45. 10.1080/15287394.2014.865581.

(70) Kavyasri, D.; Sundharesan, M.; Mathew, N. Design, Synthesis, Characterization and Insecticidal Screening of Novel Anthranilic Diamides Comprising Acyl Thiourea Substructure. Pest Manag. Sci. 2023, 79 (1), 257–273. 10.1002/ps.7196.

(71) Herath, J. M. M. K.; De Silva, W. A. P. P.; Weeraratne, T. C.; Karunaratne, S. H. P. P. Efficacy of the Insect Growth Regulator Novaluron in the Control of Dengue Vector Mosquitoes Aedes Aegypti and Ae. Albopictus. Sci. Rep. 2024, 14 (1), 1988. 10.1038/s41598-024-52384-x.

