## Supplementary material for "Instance-Wise Contrastive Graph Neural Network Enables the Discovery of Novel *Aedes aegypti* Larvicidal Compounds": Tables S1 and S2

### 1. Benchmark analysis

**Table S1. Overall predictive performance of instance-wise contrastive GNN model under the random split strategy.**

| **Task** | **Set** | **ACC ± SD** | **Recall ± SD** | **SP ± SD** | **PPV ± SD** | **NPV ± SD** | **F1 ± SD** | **G-mean ± SD** | **MCC ± SD** | **PR-AUC ± SD** | **AUC ± SD** |
| --- | --- | --- | --- | --- | --- | --- | --- | --- | --- | --- | --- |
| **0.1 µM** | Training | 0.89 ± 0.02 | 0.53 ± 0.23 | 0.95 ± 0.03 | 0.64 ± 0.05 | 0.93 ± 0.03 | 0.55 ± 0.15 | 0.69 ± 0.16 | 0.51 ± 0.14 | 0.68 ± 0.06 | 0.94 ± 0.01 |
|  | Validation | 0.93 ± 0.02 | 0.60 ± 0.38 | 0.95 ± 0.04 | 0.58 ± 0.25 | 0.97 ± 0.03 | 0.51 ± 0.18 | 0.72 ± 0.23 | 0.51 ± 0.19 | 0.60 ± 0.08 | 0.95 ± 0.02 |
|  | Test | 0.84 ± 0.02 | 0.34 ± 0.08 | 0.90 ± 0.03 | 0.34 ± 0.06 | 0.91 ± 0.01 | 0.34 ± 0.04 | 0.55 ± 0.06 | 0.25 ± 0.05 | 0.39 ± 0.06 | 0.86 ± 0.02 |
| **1 µM** | Training | 0.91 ± 0.01 | 0.96 ± 0.03 | 0.89 ± 0.03 | 0.81 ± 0.04 | 0.98 ± 0.02 | 0.88 ± 0.02 | 0.92 ± 0.01 | 0.82 ± 0.03 | 0.94 ± 0.02 | 0.98 ± 0.01 |
|  | Validation | 0.88 ± 0.02 | 0.88 ± 0.13 | 0.88 ± 0.04 | 0.68 ± 0.06 | 0.97 ± 0.04 | 0.76 ± 0.04 | 0.88 ± 0.05 | 0.70 ± 0.06 | 0.73 ± 0.11 | 0.93 ± 0.02 |
|  | Test | 0.87 ± 0.02 | 0.89 ± 0.06 | 0.86 ± 0.01 | 0.72 ± 0.01 | 0.95 ± 0.03 | 0.80 ± 0.03 | 0.87 ± 0.03 | 0.71 ± 0.05 | 0.81 ± 0.02 | 0.93 ± 0.00 |
| **10 µM** | Training | 0.95 ± 0.01 | 1.00 ± 0.00 | 0.92 ± 0.01 | 0.90 ± 0.01 | 1.00 ± 0.00 | 0.95 ± 0.01 | 0.96 ± 0.01 | 0.91 ± 0.02 | 0.97 ± 0.01 | 0.99 ± 0.01 |
|  | Validation | 0.90 ± 0.02 | 0.92 ± 0.07 | 0.89 ± 0.00 | 0.80 ± 0.01 | 0.96 ± 0.03 | 0.85 ± 0.04 | 0.91 ± 0.03 | 0.78 ± 0.05 | 0.84 ± 0.03 | 0.95 ± 0.01 |
|  | Test | 0.96 ± 0.02 | 0.99 ± 0.02 | 0.94 ± 0.01 | 0.91 ± 0.02 | 0.99 ± 0.01 | 0.95 ± 0.02 | 0.96 ± 0.02 | 0.92 ± 0.03 | 0.98 ± 0.02 | 0.99 ± 0.01 |
| **100 µM** | Training | 0.97 ± 0.01 | 0.95 ± 0.03 | 0.98 ± 0.01 | 0.98 ± 0.01 | 0.95 ± 0.03 | 0.97 ± 0.01 | 0.97 ± 0.01 | 0.93 ± 0.03 | 1.00 ± 0.00 | 1.00 ± 0.00 |
|  | Validation | 0.92 ± 0.02 | 0.90 ± 0.04 | 0.94 ± 0.05 | 0.92 ± 0.06 | 0.93 ± 0.02 | 0.90 ± 0.02 | 0.92 ± 0.02 | 0.84 ± 0.04 | 0.98 ± 0.00 | 0.98 ± 0.01 |
|  | Test | 0.86 ± 0.02 | 0.81 ± 0.04 | 0.93 ± 0.06 | 0.94 ± 0.05 | 0.80 ± 0.03 | 0.87 ± 0.02 | 0.87 ± 0.02 | 0.74 ± 0.05 | 0.96 ± 0.02 | 0.93 ± 0.04 |
| **Global** | Training | 0.93 ± 0.01 | 0.93 ± 0.03 | 0.93 ± 0.01 | 0.89 ± 0.02 | 0.96 ± 0.02 | 0.90 ± 0.02 | 0.93 ± 0.01 | 0.85 ± 0.03 | 0.97 ± 0.01 | 0.98 ± 0.00 |
|  | Validation | 0.91 ± 0.01 | 0.88 ± 0.06 | 0.92 ± 0.02 | 0.79 ± 0.04 | 0.96 ± 0.02 | 0.83 ± 0.01 | 0.90 ± 0.02 | 0.77 ± 0.02 | 0.90 ± 0.01 | 0.97 ± 0.00 |
|  | Test | 0.88 ± 0.01 | 0.84 ± 0.02 | 0.90 ± 0.01 | 0.82 ± 0.02 | 0.91 ± 0.01 | 0.83 ± 0.01 | 0.87 ± 0.01 | 0.74 ± 0.02 | 0.93 ± 0.01 | 0.95 ± 0.01 |

**Table S2. Overall predictive performance of instance-wise contrastive GNN model under the scaffold split strategy.**

| **Task** | **Set** | **ACC ± SD** | **Recall ± SD** | **SP ± SD** | **PPV ± SD** | **NPV ± SD** | **F1 ± SD** | **G-mean ± SD** | **MCC ± SD** | **PR-AUC ± SD** | **AUC** |
| --- | --- | --- | --- | --- | --- | --- | --- | --- | --- | --- | --- |
| **0.1 µM** | Training | 0.82 ± 0.06 | 0.54 ± 0.40 | 0.87 ± 0.12 | 0.53 ± 0.20 | 0.93 ± 0.05 | 0.41 ± 0.21 | 0.62 ± 0.27 | 0.38 ± 0.16 | 0.56 ± 0.09 | 0.90 ± 0.03 |
|  | Validation | 0.90 ± 0.03 | 0.20 ± 0.45 | 0.96 ± 0.05 | 0.03 ± 0.06 | 0.94 ± 0.04 | 0.04 ± 0.10 | 0.19 ± 0.42 | 0.04 ± 0.16 | 0.23 ± 0.13 | 0.78 ± 0.24 |
|  | Test | 0.86 ± 0.05 | 0.21 ± 0.31 | 0.94 ± 0.06 | 0.13 ± 0.18 | 0.91 ± 0.05 | 0.16 ± 0.21 | 0.27 ± 0.38 | 0.11 ± 0.20 | 0.36 ± 0.07 | 0.81 ± 0.08 |
| **1 µM** | Training | 0.87 ± 0.02 | 0.99 ± 0.01 | 0.80 ± 0.02 | 0.72 ± 0.03 | 0.99 ± 0.01 | 0.83 ± 0.02 | 0.89 ± 0.01 | 0.75 ± 0.03 | 0.90 ± 0.04 | 0.95 ± 0.02 |
|  | Validation | 0.87 ± 0.06 | 0.63 ± 0.20 | 0.91 ± 0.07 | 0.57 ± 0.12 | 0.93 ± 0.04 | 0.57 ± 0.11 | 0.75 ± 0.13 | 0.52 ± 0.11 | 0.62 ± 0.05 | 0.90 ± 0.08 |
|  | Test | 0.81 ± 0.07 | 0.73 ± 0.15 | 0.81 ± 0.15 | 0.57 ± 0.14 | 0.91 ± 0.04 | 0.63 ± 0.13 | 0.76 ± 0.08 | 0.51 ± 0.12 | 0.60 ± 0.14 | 0.82 ± 0.10 |
| **10 µM** | Training | 0.91 ± 0.02 | 0.98 ± 0.03 | 0.87 ± 0.02 | 0.85 ± 0.03 | 0.98 ± 0.03 | 0.91 ± 0.03 | 0.92 ± 0.02 | 0.84 ± 0.05 | 0.94 ± 0.03 | 0.96 ± 0.02 |
|  | Validation | 0.83 ± 0.09 | 0.70 ± 0.22 | 0.86 ± 0.12 | 0.64 ± 0.12 | 0.91 ± 0.08 | 0.65 ± 0.13 | 0.76 ± 0.11 | 0.55 ± 0.17 | 0.73 ± 0.16 | 0.83 ± 0.12 |
|  | Test | 0.84 ± 0.06 | 0.79 ± 0.14 | 0.85 ± 0.11 | 0.78 ± 0.12 | 0.84 ± 0.14 | 0.78 ± 0.12 | 0.81 ± 0.09 | 0.63 ± 0.16 | 0.82 ± 0.09 | 0.87 ± 0.08 |
| **100 µM** | Training | 0.94 ± 0.04 | 0.92 ± 0.07 | 0.96 ± 0.02 | 0.96 ± 0.02 | 0.91 ± 0.07 | 0.94 ± 0.04 | 0.94 ± 0.04 | 0.88 ± 0.09 | 0.98 ± 0.02 | 0.97 ± 0.03 |
|  | Validation | 0.73 ± 0.12 | 0.57 ± 0.18 | 0.82 ± 0.12 | 0.70 ± 0.13 | 0.75 ± 0.13 | 0.61 ± 0.12 | 0.67 ± 0.11 | 0.42 ± 0.21 | 0.72 ± 0.10 | 0.76 ± 0.14 |
|  | Test | 0.78 ± 0.06 | 0.69 ± 0.18 | 0.85 ± 0.07 | 0.85 ± 0.07 | 0.68 ± 0.23 | 0.75 ± 0.12 | 0.76 ± 0.09 | 0.53 ± 0.15 | 0.86 ± 0.08 | 0.82 ± 0.09 |
| **Global** | Training | 0.89 ± 0.04 | 0.92 ± 0.04 | 0.87 ± 0.05 | 0.80 ± 0.07 | 0.95 ± 0.02 | 0.85 ± 0.04 | 0.89 ± 0.03 | 0.77 ± 0.07 | 0.94 ± 0.03 | 0.96 ± 0.02 |
|  | Validation | 0.83 ± 0.06 | 0.58 ± 0.12 | 0.90 ± 0.05 | 0.60 ± 0.09 | 0.89 ± 0.05 | 0.58 ± 0.09 | 0.71 ± 0.07 | 0.48 ± 0.12 | 0.64 ± 0.09 | 0.83 ± 0.10 |
|  | Test | 0.82 ± 0.05 | 0.67 ± 0.15 | 0.87 ± 0.08 | 0.72 ± 0.06 | 0.85 ± 0.08 | 0.69 ± 0.11 | 0.76 ± 0.08 | 0.56 ± 0.11 | 0.75 ± 0.08 | 0.85 ± 0.06 |

**Table S3. Overall predictive performance of the uncertainty-weighted instance-wise contrastive GNN under the random split strategy.**

| **Task** | **Set** | **ACC ± SD** | **Recall ± SD** | **SP ± SD** | **PPV ± SD** | **NPV ± SD** | **F1 ± SD** | **G-mean ± SD** | **MCC ± SD** | **PR-AUC ± SD** | **AUC** |
| --- | --- | --- | --- | --- | --- | --- | --- | --- | --- | --- | --- |
| **0.1 µM** | Training | 0.89 ± 0.03 | 0.40 ± 0.31 | 0.97 ± 0.02 | 0.72 ± 0.10 | 0.91 ± 0.04 | 0.46 ± 0.27 | 0.58 ± 0.26 | 0.46 ± 0.23 | 0.74 ± 0.08 | 0.95 ± 0.01 |
|  | Validation | 0.93 ± 0.01 | 0.30 ± 0.11 | 0.98 ± 0.02 | 0.70 ± 0.27 | 0.95 ± 0.01 | 0.39 ± 0.07 | 0.54 ± 0.09 | 0.41 ± 0.09 | 0.68 ± 0.08 | 0.94 ± 0.04 |
|  | Test | 0.89 ± 0.03 | 0.31 ± 0.23 | 0.97 ± 0.05 | 0.59 ± 0.43 | 0.91 ± 0.03 | 0.36 ± 0.25 | 0.48 ± 0.30 | 0.36 ± 0.24 | 0.54 ± 0.11 | 0.89 ± 0.02 |
| **1 µM** | Training | 0.93 ± 0.02 | 0.95 ± 0.04 | 0.92 ± 0.03 | 0.86 ± 0.05 | 0.98 ± 0.02 | 0.90 ± 0.02 | 0.94 ± 0.01 | 0.85 ± 0.03 | 0.96 ± 0.02 | 0.98 ± 0.01 |
|  | Validation | 0.88 ± 0.06 | 0.87 ± 0.13 | 0.88 ± 0.05 | 0.67 ± 0.11 | 0.96 ± 0.04 | 0.76 ± 0.11 | 0.87 ± 0.08 | 0.69 ± 0.15 | 0.79 ± 0.05 | 0.94 ± 0.02 |
|  | Test | 0.86 ± 0.02 | 0.87 ± 0.05 | 0.86 ± 0.02 | 0.73 ± 0.03 | 0.94 ± 0.02 | 0.79 ± 0.03 | 0.87 ± 0.03 | 0.70 ± 0.05 | 0.87 ± 0.04 | 0.95 ± 0.02 |
| **10 µM** | Training | 0.95 ± 0.01 | 0.98 ± 0.01 | 0.93 ± 0.02 | 0.92 ± 0.02 | 0.99 ± 0.01 | 0.95 ± 0.01 | 0.96 ± 0.01 | 0.91 ± 0.02 | 0.98 ± 0.01 | 0.99 ± 0.01 |
|  | Validation | 0.91 ± 0.02 | 0.93 ± 0.06 | 0.90 ± 0.01 | 0.81 ± 0.02 | 0.97 ± 0.03 | 0.86 ± 0.04 | 0.91 ± 0.03 | 0.80 ± 0.06 | 0.84 ± 0.04 | 0.95 ± 0.01 |
|  | Test | 0.94 ± 0.01 | 0.96 ± 0.03 | 0.94 ± 0.02 | 0.91 ± 0.03 | 0.97 ± 0.02 | 0.93 ± 0.02 | 0.95 ± 0.01 | 0.89 ± 0.03 | 0.97 ± 0.01 | 0.98 ± 0.01 |
| **100 µM** | Training | 0.98 ± 0.01 | 0.98 ± 0.02 | 0.98 ± 0.02 | 0.99 ± 0.02 | 0.97 ± 0.03 | 0.98 ± 0.01 | 0.98 ± 0.01 | 0.96 ± 0.03 | 1.00 ± 0.00 | 1.00 ± 0.00 |
|  | Validation | 0.93 ± 0.03 | 0.93 ± 0.04 | 0.93 ± 0.05 | 0.90 ± 0.06 | 0.95 ± 0.02 | 0.92 ± 0.03 | 0.93 ± 0.02 | 0.85 ± 0.05 | 0.98 ± 0.01 | 0.98 ± 0.01 |
|  | Test | 0.89 ± 0.02 | 0.84 ± 0.06 | 0.96 ± 0.05 | 0.97 ± 0.04 | 0.83 ± 0.05 | 0.90 ± 0.02 | 0.90 ± 0.02 | 0.80 ± 0.04 | 0.96 ± 0.02 | 0.93 ± 0.03 |
| **Global** | Training | 0.94 ± 0.02 | 0.91 ± 0.04 | 0.95 ± 0.01 | 0.92 ± 0.02 | 0.95 ± 0.02 | 0.92 ± 0.02 | 0.93 ± 0.02 | 0.87 ± 0.03 | 0.98 ± 0.01 | 0.99 ± 0.01 |
|  | Validation | 0.91 ± 0.02 | 0.87 ± 0.05 | 0.93 ± 0.02 | 0.80 ± 0.04 | 0.95 ± 0.02 | 0.83 ± 0.04 | 0.90 ± 0.03 | 0.78 ± 0.05 | 0.91 ± 0.02 | 0.97 ± 0.01 |
|  | Test | 0.90 ± 0.01 | 0.84 ± 0.03 | 0.93 ± 0.02 | 0.86 ± 0.03 | 0.91 ± 0.02 | 0.85 ± 0.02 | 0.88 ± 0.02 | 0.77 ± 0.03 | 0.93 ± 0.01 | 0.95 ± 0.01 |

**Table S4. Overall predictive performance of the SVM under the random split strategy.**

| **Task** | **Set** | **ACC ± SD** | **Recall ± SD** | **SP ± SD** | **PPV ± SD** | **NPV ± SD** | **F1 ± SD** | **G-mean ± SD** | **MCC ± SD** | **PR-AUC ± SD** | **AUC** |
| --- | --- | --- | --- | --- | --- | --- | --- | --- | --- | --- | --- |
| **0.1 µM** | Training | 0.88 ± 0.01 | 0.10 ± 0.06 | 1.00 ± 0.00 | 1.00 ± 0.00 | 0.88 ± 0.01 | 0.17 ± 0.10 | 0.29 ± 0.11 | 0.27 ± 0.10 | 0.69 ± 0.07 | 0.82 ± 0.05 |
|  | Validation | 0.86 ± 0.00 | 0.01 ± 0.01 | 0.99 ± 0.00 | 0.08 ± 0.11 | 0.87 ± 0.00 | 0.02 ± 0.02 | 0.06 ± 0.08 | -0.01 ± 0.04 | 0.18 ± 0.03 | 0.53 ± 0.04 |
|  | Test | 0.88 ± 0.01 | 0.03 ± 0.06 | 1.00 ± 0.01 | 0.20 ± 0.45 | 0.88 ± 0.01 | 0.05 ± 0.11 | 0.08 ± 0.17 | 0.06 ± 0.17 | 0.40 ± 0.06 | 0.85 ± 0.06 |
| **1 µM** | Training | 0.61 ± 0.08 | 0.89 ± 0.05 | 0.49 ± 0.14 | 0.45 ± 0.07 | 0.91 ± 0.03 | 0.59 ± 0.05 | 0.65 ± 0.07 | 0.37 ± 0.09 | 0.85 ± 0.03 | 0.88 ± 0.03 |
|  | Validation | 0.57 ± 0.05 | 0.53 ± 0.07 | 0.59 ± 0.08 | 0.37 ± 0.04 | 0.74 ± 0.02 | 0.43 ± 0.04 | 0.56 ± 0.04 | 0.11 ± 0.06 | 0.46 ± 0.02 | 0.64 ± 0.01 |
|  | Test | 0.50 ± 0.19 | 0.95 ± 0.11 | 0.31 ± 0.32 | 0.42 ± 0.17 | 0.98 ± 0.05 | 0.55 ± 0.11 | 0.49 ± 0.19 | 0.32 ± 0.18 | 0.77 ± 0.03 | 0.92 ± 0.01 |
| **10 µM** | Training | 0.58 ± 0.03 | 1.00 ± 0.00 | 0.29 ± 0.06 | 0.50 ± 0.02 | 1.00 ± 0.00 | 0.67 ± 0.02 | 0.54 ± 0.05 | 0.38 ± 0.05 | 0.95 ± 0.03 | 0.96 ± 0.03 |
|  | Validation | 0.54 ± 0.03 | 0.93 ± 0.12 | 0.27 ± 0.12 | 0.47 ± 0.02 | 0.89 ± 0.11 | 0.62 ± 0.03 | 0.48 ± 0.07 | 0.26 ± 0.07 | 0.68 ± 0.06 | 0.74 ± 0.07 |
|  | Test | 0.47 ± 0.02 | 1.00 ± 0.00 | 0.12 ± 0.04 | 0.43 ± 0.01 | 1.00 ± 0.00 | 0.61 ± 0.01 | 0.34 ± 0.06 | 0.22 ± 0.04 | 0.83 ± 0.02 | 0.92 ± 0.02 |
| **100 µM** | Training | 0.60 ± 0.01 | 1.00 ± 0.00 | 0.16 ± 0.02 | 0.57 ± 0.01 | 1.00 ± 0.00 | 0.73 ± 0.01 | 0.40 ± 0.03 | 0.31 ± 0.02 | 0.92 ± 0.02 | 0.90 ± 0.02 |
|  | Validation | 0.52 ± 0.01 | 0.77 ± 0.01 | 0.25 ± 0.01 | 0.53 ± 0.01 | 0.49 ± 0.02 | 0.63 ± 0.01 | 0.44 ± 0.01 | 0.02 ± 0.02 | 0.67 ± 0.03 | 0.58 ± 0.03 |
|  | Test | 0.59 ± 0.01 | 1.00 ± 0.00 | 0.06 ± 0.02 | 0.58 ± 0.01 | 1.00 ± 0.00 | 0.73 ± 0.00 | 0.25 ± 0.05 | 0.19 ± 0.04 | 0.94 ± 0.01 | 0.91 ± 0.02 |
| **Global** | Training | 0.67 ± 0.02 | 0.89 ± 0.01 | 0.56 ± 0.03 | 0.51 ± 0.02 | 0.90 ± 0.01 | 0.65 ± 0.01 | 0.70 ± 0.02 | 0.43 ± 0.02 | 0.75 ± 0.03 | 0.82 ± 0.03 |
|  | Validation | 0.62 ± 0.02 | 0.69 ± 0.05 | 0.59 ± 0.04 | 0.47 ± 0.02 | 0.78 ± 0.02 | 0.56 ± 0.02 | 0.64 ± 0.02 | 0.27 ± 0.03 | 0.52 ± 0.02 | 0.69 ± 0.01 |
|  | Test | 0.61 ± 0.04 | 0.90 ± 0.02 | 0.46 ± 0.08 | 0.47 ± 0.03 | 0.90 ± 0.01 | 0.62 ± 0.02 | 0.64 ± 0.04 | 0.36 ± 0.04 | 0.63 ± 0.03 | 0.77 ± 0.03 |

**Table S5. Overall predictive performance of the Random Forest under the random split strategy.**

| **Task** | **Set** | **ACC ± SD** | **Recall ± SD** | **SP ± SD** | **PPV ± SD** | **NPV ± SD** | **F1 ± SD** | **G-mean ± SD** | **MCC ± SD** | **PR-AUC ± SD** | **AUC** |
| --- | --- | --- | --- | --- | --- | --- | --- | --- | --- | --- | --- |
| **0.1 µM** | Training | 0.87 ± 0.00 | 0.00 ± 0.00 | 1.00 ± 0.00 | 0.00 ± 0.00 | 0.87 ± 0.00 | 0.00 ± 0.00 | 0.00 ± 0.00 | 0.00 ± 0.00 | 0.54 ± 0.02 | 0.87 ± 0.00 |
|  | Validation | 0.87 ± 0.00 | 0.00 ± 0.00 | 1.00 ± 0.00 | 0.00 ± 0.00 | 0.87 ± 0.00 | 0.00 ± 0.00 | 0.00 ± 0.00 | 0.00 ± 0.00 | 0.47 ± 0.04 | 0.83 ± 0.02 |
|  | Test | 0.88 ± 0.00 | 0.00 ± 0.00 | 1.00 ± 0.00 | 0.00 ± 0.00 | 0.88 ± 0.00 | 0.00 ± 0.00 | 0.00 ± 0.00 | 0.00 ± 0.00 | 0.58 ± 0.07 | 0.88 ± 0.02 |
| **1 µM** | Training | 0.83 ± 0.00 | 0.65 ± 0.01 | 0.91 ± 0.00 | 0.77 ± 0.00 | 0.86 ± 0.00 | 0.71 ± 0.01 | 0.77 ± 0.01 | 0.60 ± 0.01 | 0.81 ± 0.00 | 0.91 ± 0.00 |
|  | Validation | 0.83 ± 0.01 | 0.64 ± 0.02 | 0.91 ± 0.00 | 0.77 ± 0.01 | 0.85 ± 0.01 | 0.70 ± 0.01 | 0.77 ± 0.01 | 0.59 ± 0.01 | 0.80 ± 0.01 | 0.89 ± 0.00 |
|  | Test | 0.81 ± 0.03 | 0.54 ± 0.06 | 0.93 ± 0.02 | 0.77 ± 0.07 | 0.83 ± 0.02 | 0.63 ± 0.07 | 0.71 ± 0.05 | 0.53 ± 0.09 | 0.79 ± 0.04 | 0.87 ± 0.03 |
| **10 µM** | Training | 0.80 ± 0.00 | 0.62 ± 0.01 | 0.93 ± 0.00 | 0.85 ± 0.00 | 0.77 ± 0.01 | 0.72 ± 0.01 | 0.76 ± 0.01 | 0.58 ± 0.01 | 0.87 ± 0.00 | 0.89 ± 0.00 |
|  | Validation | 0.80 ± 0.01 | 0.62 ± 0.02 | 0.93 ± 0.00 | 0.85 ± 0.01 | 0.78 ± 0.01 | 0.72 ± 0.01 | 0.76 ± 0.01 | 0.59 ± 0.01 | 0.85 ± 0.01 | 0.87 ± 0.01 |
|  | Test | 0.74 ± 0.01 | 0.44 ± 0.02 | 0.95 ± 0.01 | 0.85 ± 0.03 | 0.72 ± 0.01 | 0.58 ± 0.02 | 0.65 ± 0.01 | 0.47 ± 0.02 | 0.85 ± 0.01 | 0.89 ± 0.01 |
| **100 µM** | Training | 0.81 ± 0.01 | 0.66 ± 0.01 | 0.99 ± 0.00 | 0.98 ± 0.00 | 0.72 ± 0.01 | 0.79 ± 0.01 | 0.81 ± 0.01 | 0.67 ± 0.01 | 0.92 ± 0.00 | 0.89 ± 0.00 |
|  | Validation | 0.81 ± 0.01 | 0.66 ± 0.02 | 0.97 ± 0.01 | 0.97 ± 0.01 | 0.72 ± 0.01 | 0.78 ± 0.01 | 0.80 ± 0.01 | 0.66 ± 0.01 | 0.91 ± 0.00 | 0.88 ± 0.01 |
|  | Test | 0.74 ± 0.02 | 0.58 ± 0.03 | 0.96 ± 0.00 | 0.95 ± 0.00 | 0.64 ± 0.01 | 0.72 ± 0.02 | 0.74 ± 0.02 | 0.56 ± 0.02 | 0.88 ± 0.01 | 0.81 ± 0.02 |
| **Global** | Training | 0.83 ± 0.00 | 0.58 ± 0.01 | 0.96 ± 0.00 | 0.88 ± 0.00 | 0.81 ± 0.00 | 0.70 ± 0.00 | 0.75 ± 0.00 | 0.61 ± 0.00 | 0.85 ± 0.00 | 0.89 ± 0.00 |
|  | Validation | 0.83 ± 0.00 | 0.58 ± 0.01 | 0.96 ± 0.00 | 0.87 ± 0.00 | 0.81 ± 0.00 | 0.70 ± 0.01 | 0.74 ± 0.01 | 0.61 ± 0.01 | 0.84 ± 0.00 | 0.88 ± 0.00 |
|  | Test | 0.79 ± 0.01 | 0.48 ± 0.03 | 0.96 ± 0.01 | 0.87 ± 0.03 | 0.78 ± 0.01 | 0.62 ± 0.02 | 0.68 ± 0.02 | 0.53 ± 0.03 | 0.81 ± 0.02 | 0.87 ± 0.01 |

**Table S6. Overall predictive performance of the MPNN under the random split strategy.**

| **Task** | **Set** | **ACC ± SD** | **Recall ± SD** | **SP ± SD** | **PPV ± SD** | **NPV ± SD** | **F1 ± SD** | **G-mean ± SD** | **MCC ± SD** | **PR-AUC ± SD** | **AUC** |
| --- | --- | --- | --- | --- | --- | --- | --- | --- | --- | --- | --- |
| **0.1 µM** | Training | 0.85 ± 0.01 | 0.11 ± 0.25 | 0.98 ± 0.05 | 0.09 ± 0.20 | 0.87 ± 0.03 | 0.10 ± 0.22 | 0.14 ± 0.31 | 0.07 ± 0.19 | 0.38 ± 0.03 | 0.84 ± 0.02 |
|  | Validation | 0.91 ± 0.04 | 0.10 ± 0.22 | 0.97 ± 0.06 | 0.04 ± 0.10 | 0.93 ± 0.01 | 0.06 ± 0.14 | 0.13 ± 0.29 | 0.04 ± 0.12 | 0.27 ± 0.09 | 0.77 ± 0.07 |
|  | Test | 0.87 ± 0.02 | 0.11 ± 0.26 | 0.97 ± 0.06 | 0.07 ± 0.16 | 0.89 ± 0.03 | 0.09 ± 0.20 | 0.14 ± 0.31 | 0.07 ± 0.16 | 0.43 ± 0.12 | 0.84 ± 0.04 |
| **1 µM** | Training | 0.84 ± 0.02 | 0.73 ± 0.14 | 0.88 ± 0.03 | 0.75 ± 0.02 | 0.88 ± 0.05 | 0.73 ± 0.07 | 0.80 ± 0.07 | 0.62 ± 0.07 | 0.76 ± 0.02 | 0.90 ± 0.01 |
|  | Validation | 0.84 ± 0.04 | 0.75 ± 0.08 | 0.86 ± 0.03 | 0.60 ± 0.07 | 0.93 ± 0.02 | 0.67 ± 0.07 | 0.80 ± 0.05 | 0.57 ± 0.10 | 0.64 ± 0.14 | 0.89 ± 0.03 |
|  | Test | 0.80 ± 0.04 | 0.64 ± 0.10 | 0.87 ± 0.08 | 0.69 ± 0.11 | 0.85 ± 0.02 | 0.65 ± 0.05 | 0.74 ± 0.04 | 0.52 ± 0.06 | 0.72 ± 0.09 | 0.86 ± 0.04 |
| **10 µM** | Training | 0.83 ± 0.02 | 0.72 ± 0.06 | 0.90 ± 0.02 | 0.85 ± 0.02 | 0.82 ± 0.03 | 0.78 ± 0.03 | 0.81 ± 0.03 | 0.64 ± 0.03 | 0.85 ± 0.03 | 0.89 ± 0.02 |
|  | Validation | 0.87 ± 0.02 | 0.76 ± 0.04 | 0.92 ± 0.02 | 0.81 ± 0.04 | 0.90 ± 0.02 | 0.79 ± 0.03 | 0.84 ± 0.02 | 0.70 ± 0.04 | 0.82 ± 0.06 | 0.92 ± 0.03 |
|  | Test | 0.79 ± 0.04 | 0.63 ± 0.10 | 0.90 ± 0.05 | 0.81 ± 0.08 | 0.78 ± 0.04 | 0.70 ± 0.07 | 0.75 ± 0.06 | 0.56 ± 0.09 | 0.82 ± 0.05 | 0.86 ± 0.03 |
| **100 µM** | Training | 0.82 ± 0.01 | 0.70 ± 0.04 | 0.96 ± 0.02 | 0.96 ± 0.02 | 0.73 ± 0.02 | 0.81 ± 0.02 | 0.82 ± 0.02 | 0.68 ± 0.02 | 0.92 ± 0.02 | 0.89 ± 0.02 |
|  | Validation | 0.89 ± 0.01 | 0.75 ± 0.05 | 0.99 ± 0.01 | 0.99 ± 0.02 | 0.85 ± 0.02 | 0.85 ± 0.02 | 0.86 ± 0.02 | 0.79 ± 0.02 | 0.93 ± 0.02 | 0.92 ± 0.03 |
|  | Test | 0.77 ± 0.05 | 0.63 ± 0.10 | 0.95 ± 0.03 | 0.95 ± 0.04 | 0.67 ± 0.06 | 0.75 ± 0.07 | 0.77 ± 0.06 | 0.60 ± 0.08 | 0.88 ± 0.04 | 0.81 ± 0.04 |
| **Global** | Training | 0.83 ± 0.01 | 0.66 ± 0.07 | 0.93 ± 0.03 | 0.85 ± 0.04 | 0.83 ± 0.03 | 0.74 ± 0.03 | 0.78 ± 0.03 | 0.63 ± 0.02 | 0.85 ± 0.02 | 0.90 ± 0.02 |
|  | Validation | 0.88 ± 0.01 | 0.71 ± 0.04 | 0.93 ± 0.02 | 0.79 ± 0.04 | 0.90 ± 0.01 | 0.74 ± 0.01 | 0.81 ± 0.02 | 0.67 ± 0.02 | 0.81 ± 0.04 | 0.91 ± 0.02 |
|  | Test | 0.81 ± 0.03 | 0.58 ± 0.09 | 0.92 ± 0.05 | 0.82 ± 0.08 | 0.81 ± 0.03 | 0.67 ± 0.05 | 0.73 ± 0.05 | 0.56 ± 0.06 | 0.80 ± 0.03 | 0.86 ± 0.03 |

**Table S7. Overall predictive performance of the GAT under the random split strategy.**

| **Task** | **Set** | **ACC ± SD** | **Recall ± SD** | **SP ± SD** | **PPV ± SD** | **NPV ± SD** | **F1 ± SD** | **G-mean ± SD** | **MCC ± SD** | **PR-AUC ± SD** | **AUC** |
| --- | --- | --- | --- | --- | --- | --- | --- | --- | --- | --- | --- |
| **0.1 µM** | Training | 0.86 ± 0.00 | 0.00 ± 0.00 | 1.00 ± 0.00 | 0.00 ± 0.00 | 0.86 ± 0.00 | 0.00 ± 0.00 | 0.00 ± 0.00 | 0.00 ± 0.00 | 0.33 ± 0.03 | 0.81 ± 0.01 |
|  | Validation | 0.93 ± 0.00 | 0.00 ± 0.00 | 1.00 ± 0.00 | 0.00 ± 0.00 | 0.93 ± 0.00 | 0.00 ± 0.00 | 0.00 ± 0.00 | 0.00 ± 0.00 | 0.15 ± 0.02 | 0.71 ± 0.02 |
|  | Test | 0.88 ± 0.00 | 0.00 ± 0.00 | 1.00 ± 0.00 | 0.00 ± 0.00 | 0.88 ± 0.00 | 0.00 ± 0.00 | 0.00 ± 0.00 | 0.00 ± 0.00 | 0.41 ± 0.10 | 0.85 ± 0.02 |
| **1 µM** | Training | 0.84 ± 0.01 | 0.89 ± 0.05 | 0.82 ± 0.02 | 0.70 ± 0.02 | 0.94 ± 0.02 | 0.79 ± 0.02 | 0.86 ± 0.02 | 0.68 ± 0.03 | 0.76 ± 0.04 | 0.90 ± 0.01 |
|  | Validation | 0.84 ± 0.02 | 0.90 ± 0.04 | 0.83 ± 0.03 | 0.60 ± 0.04 | 0.97 ± 0.01 | 0.72 ± 0.03 | 0.86 ± 0.02 | 0.64 ± 0.04 | 0.62 ± 0.07 | 0.89 ± 0.02 |
|  | Test | 0.82 ± 0.02 | 0.85 ± 0.09 | 0.81 ± 0.01 | 0.65 ± 0.01 | 0.93 ± 0.04 | 0.74 ± 0.04 | 0.83 ± 0.04 | 0.62 ± 0.06 | 0.70 ± 0.08 | 0.87 ± 0.02 |
| **10 µM** | Training | 0.85 ± 0.01 | 0.83 ± 0.03 | 0.87 ± 0.02 | 0.82 ± 0.01 | 0.87 ± 0.02 | 0.83 ± 0.01 | 0.85 ± 0.01 | 0.70 ± 0.02 | 0.84 ± 0.02 | 0.88 ± 0.01 |
|  | Validation | 0.90 ± 0.02 | 0.88 ± 0.04 | 0.91 ± 0.02 | 0.82 ± 0.04 | 0.95 ± 0.02 | 0.85 ± 0.03 | 0.90 ± 0.02 | 0.78 ± 0.05 | 0.77 ± 0.09 | 0.90 ± 0.05 |
|  | Test | 0.84 ± 0.03 | 0.82 ± 0.05 | 0.85 ± 0.04 | 0.79 ± 0.04 | 0.87 ± 0.03 | 0.80 ± 0.03 | 0.83 ± 0.03 | 0.67 ± 0.06 | 0.84 ± 0.03 | 0.90 ± 0.02 |
| **100 µM** | Training | 0.84 ± 0.01 | 0.76 ± 0.02 | 0.94 ± 0.03 | 0.93 ± 0.03 | 0.77 ± 0.01 | 0.84 ± 0.01 | 0.84 ± 0.01 | 0.70 ± 0.02 | 0.92 ± 0.01 | 0.88 ± 0.01 |
|  | Validation | 0.90 ± 0.02 | 0.78 ± 0.03 | 0.99 ± 0.03 | 0.98 ± 0.04 | 0.86 ± 0.02 | 0.87 ± 0.03 | 0.88 ± 0.02 | 0.81 ± 0.04 | 0.93 ± 0.02 | 0.92 ± 0.03 |
|  | Test | 0.80 ± 0.02 | 0.71 ± 0.02 | 0.91 ± 0.03 | 0.91 ± 0.03 | 0.71 ± 0.02 | 0.80 ± 0.02 | 0.80 ± 0.02 | 0.62 ± 0.04 | 0.92 ± 0.01 | 0.89 ± 0.02 |
| **Global** | Training | 0.85 ± 0.01 | 0.73 ± 0.03 | 0.91 ± 0.01 | 0.82 ± 0.02 | 0.86 ± 0.01 | 0.78 ± 0.01 | 0.82 ± 0.01 | 0.66 ± 0.02 | 0.86 ± 0.01 | 0.90 ± 0.01 |
|  | Validation | 0.89 ± 0.01 | 0.78 ± 0.03 | 0.93 ± 0.02 | 0.80 ± 0.04 | 0.93 ± 0.01 | 0.79 ± 0.02 | 0.85 ± 0.01 | 0.72 ± 0.03 | 0.85 ± 0.03 | 0.91 ± 0.01 |
|  | Test | 0.83 ± 0.01 | 0.71 ± 0.03 | 0.90 ± 0.02 | 0.79 ± 0.03 | 0.85 ± 0.01 | 0.75 ± 0.02 | 0.80 ± 0.02 | 0.63 ± 0.03 | 0.84 ± 0.01 | 0.90 ± 0.01 |

**Table S8. Overall predictive performance of the GIN under the random split strategy.**

| **Task** | **Set** | **ACC ± SD** | **Recall ± SD** | **SP ± SD** | **PPV ± SD** | **NPV ± SD** | **F1 ± SD** | **G-mean ± SD** | **MCC ± SD** | **PR-AUC ± SD** | **AUC** |
| --- | --- | --- | --- | --- | --- | --- | --- | --- | --- | --- | --- |
| **0.1 µM** | Training | 0.86 ± 0.00 | 0.00 ± 0.00 | 1.00 ± 0.00 | 0.00 ± 0.00 | 0.86 ± 0.00 | 0.00 ± 0.00 | 0.00 ± 0.00 | 0.00 ± 0.00 | 0.35 ± 0.04 | 0.82 ± 0.02 |
|  | Validation | 0.93 ± 0.00 | 0.00 ± 0.00 | 1.00 ± 0.00 | 0.00 ± 0.00 | 0.93 ± 0.00 | 0.00 ± 0.00 | 0.00 ± 0.00 | 0.00 ± 0.00 | 0.24 ± 0.09 | 0.78 ± 0.07 |
|  | Test | 0.88 ± 0.00 | 0.00 ± 0.00 | 1.00 ± 0.00 | 0.00 ± 0.00 | 0.88 ± 0.00 | 0.00 ± 0.00 | 0.00 ± 0.00 | 0.00 ± 0.00 | 0.47 ± 0.16 | 0.82 ± 0.03 |
| **1 µM** | Training | 0.84 ± 0.02 | 0.80 ± 0.06 | 0.85 ± 0.04 | 0.72 ± 0.04 | 0.90 ± 0.03 | 0.76 ± 0.03 | 0.82 ± 0.02 | 0.64 ± 0.04 | 0.75 ± 0.03 | 0.90 ± 0.01 |
|  | Validation | 0.83 ± 0.02 | 0.82 ± 0.04 | 0.84 ± 0.02 | 0.58 ± 0.03 | 0.94 ± 0.01 | 0.68 ± 0.02 | 0.83 ± 0.02 | 0.59 ± 0.03 | 0.70 ± 0.06 | 0.91 ± 0.00 |
|  | Test | 0.78 ± 0.04 | 0.67 ± 0.12 | 0.83 ± 0.08 | 0.64 ± 0.07 | 0.86 ± 0.04 | 0.65 ± 0.06 | 0.74 ± 0.05 | 0.50 ± 0.07 | 0.72 ± 0.07 | 0.85 ± 0.04 |
| **10 µM** | Training | 0.84 ± 0.01 | 0.79 ± 0.07 | 0.88 ± 0.04 | 0.83 ± 0.03 | 0.85 ± 0.04 | 0.81 ± 0.02 | 0.83 ± 0.02 | 0.68 ± 0.03 | 0.85 ± 0.03 | 0.90 ± 0.01 |
|  | Validation | 0.88 ± 0.03 | 0.86 ± 0.07 | 0.89 ± 0.03 | 0.78 ± 0.05 | 0.93 ± 0.03 | 0.82 ± 0.05 | 0.87 ± 0.04 | 0.73 ± 0.07 | 0.82 ± 0.08 | 0.91 ± 0.04 |
|  | Test | 0.81 ± 0.05 | 0.75 ± 0.14 | 0.86 ± 0.09 | 0.79 ± 0.07 | 0.84 ± 0.06 | 0.76 ± 0.07 | 0.80 ± 0.06 | 0.62 ± 0.09 | 0.83 ± 0.03 | 0.87 ± 0.04 |
| **100 µM** | Training | 0.83 ± 0.02 | 0.75 ± 0.06 | 0.93 ± 0.04 | 0.93 ± 0.03 | 0.76 ± 0.04 | 0.83 ± 0.02 | 0.83 ± 0.02 | 0.68 ± 0.02 | 0.92 ± 0.01 | 0.90 ± 0.02 |
|  | Validation | 0.87 ± 0.03 | 0.77 ± 0.05 | 0.94 ± 0.05 | 0.91 ± 0.07 | 0.85 ± 0.03 | 0.83 ± 0.04 | 0.85 ± 0.03 | 0.73 ± 0.07 | 0.90 ± 0.07 | 0.91 ± 0.05 |
|  | Test | 0.79 ± 0.04 | 0.68 ± 0.10 | 0.93 ± 0.05 | 0.93 ± 0.04 | 0.70 ± 0.06 | 0.78 ± 0.06 | 0.79 ± 0.05 | 0.62 ± 0.06 | 0.91 ± 0.04 | 0.88 ± 0.05 |
| **Global** | Training | 0.84 ± 0.01 | 0.70 ± 0.05 | 0.92 ± 0.02 | 0.83 ± 0.03 | 0.85 ± 0.02 | 0.76 ± 0.02 | 0.80 ± 0.02 | 0.65 ± 0.01 | 0.86 ± 0.01 | 0.91 ± 0.01 |
|  | Validation | 0.88 ± 0.02 | 0.75 ± 0.05 | 0.92 ± 0.02 | 0.77 ± 0.05 | 0.92 ± 0.01 | 0.76 ± 0.04 | 0.83 ± 0.03 | 0.68 ± 0.05 | 0.83 ± 0.06 | 0.92 ± 0.03 |
|  | Test | 0.82 ± 0.02 | 0.64 ± 0.10 | 0.91 ± 0.05 | 0.80 ± 0.06 | 0.83 ± 0.03 | 0.70 ± 0.05 | 0.76 ± 0.05 | 0.59 ± 0.05 | 0.81 ± 0.04 | 0.88 ± 0.03 |

**Table S9. Overall predictive performance of the AttentiveFP under the random split strategy.**

| **Task** | **Set** | **ACC ± SD** | **Recall ± SD** | **SP ± SD** | **PPV ± SD** | **NPV ± SD** | **F1 ± SD** | **G-mean ± SD** | **MCC ± SD** | **PR-AUC ± SD** | **AUC** |
| --- | --- | --- | --- | --- | --- | --- | --- | --- | --- | --- | --- |
| **0.1 µM** | Training | 0.86 ± 0.00 | 0.00 ± 0.00 | 1.00 ± 0.00 | 0.00 ± 0.00 | 0.86 ± 0.00 | 0.00 ± 0.00 | 0.00 ± 0.00 | 0.00 ± 0.00 | 0.37 ± 0.02 | 0.84 ± 0.01 |
|  | Validation | 0.93 ± 0.00 | 0.00 ± 0.00 | 1.00 ± 0.00 | 0.00 ± 0.00 | 0.93 ± 0.00 | 0.00 ± 0.00 | 0.00 ± 0.00 | 0.00 ± 0.00 | 0.22 ± 0.03 | 0.83 ± 0.03 |
|  | Test | 0.88 ± 0.00 | 0.00 ± 0.00 | 1.00 ± 0.00 | 0.00 ± 0.00 | 0.88 ± 0.00 | 0.00 ± 0.00 | 0.00 ± 0.00 | 0.00 ± 0.00 | 0.34 ± 0.08 | 0.81 ± 0.04 |
| **1 µM** | Training | 0.86 ± 0.00 | 0.94 ± 0.03 | 0.83 ± 0.02 | 0.72 ± 0.01 | 0.97 ± 0.01 | 0.82 ± 0.01 | 0.88 ± 0.01 | 0.73 ± 0.01 | 0.73 ± 0.02 | 0.91 ± 0.00 |
|  | Validation | 0.87 ± 0.01 | 0.93 ± 0.07 | 0.85 ± 0.03 | 0.63 ± 0.03 | 0.98 ± 0.02 | 0.75 ± 0.02 | 0.89 ± 0.02 | 0.69 ± 0.03 | 0.59 ± 0.04 | 0.91 ± 0.01 |
|  | Test | 0.79 ± 0.01 | 0.85 ± 0.05 | 0.77 ± 0.03 | 0.61 ± 0.02 | 0.92 ± 0.02 | 0.71 ± 0.02 | 0.80 ± 0.02 | 0.57 ± 0.03 | 0.68 ± 0.05 | 0.87 ± 0.02 |
| **10 µM** | Training | 0.88 ± 0.01 | 0.87 ± 0.02 | 0.88 ± 0.01 | 0.85 ± 0.01 | 0.90 ± 0.01 | 0.86 ± 0.01 | 0.88 ± 0.01 | 0.75 ± 0.02 | 0.83 ± 0.02 | 0.90 ± 0.01 |
|  | Validation | 0.92 ± 0.02 | 0.93 ± 0.08 | 0.92 ± 0.02 | 0.84 ± 0.03 | 0.97 ± 0.03 | 0.88 ± 0.04 | 0.92 ± 0.03 | 0.83 ± 0.05 | 0.86 ± 0.04 | 0.94 ± 0.01 |
|  | Test | 0.84 ± 0.02 | 0.84 ± 0.06 | 0.84 ± 0.02 | 0.78 ± 0.02 | 0.89 ± 0.04 | 0.81 ± 0.03 | 0.84 ± 0.03 | 0.67 ± 0.05 | 0.79 ± 0.06 | 0.87 ± 0.03 |
| **100 µM** | Training | 0.87 ± 0.01 | 0.80 ± 0.01 | 0.96 ± 0.01 | 0.96 ± 0.01 | 0.80 ± 0.01 | 0.87 ± 0.01 | 0.87 ± 0.01 | 0.75 ± 0.01 | 0.91 ± 0.01 | 0.90 ± 0.01 |
|  | Validation | 0.91 ± 0.02 | 0.83 ± 0.04 | 0.96 ± 0.03 | 0.94 ± 0.04 | 0.89 ± 0.03 | 0.88 ± 0.03 | 0.89 ± 0.03 | 0.81 ± 0.05 | 0.95 ± 0.02 | 0.94 ± 0.02 |
|  | Test | 0.86 ± 0.02 | 0.78 ± 0.04 | 0.96 ± 0.00 | 0.96 ± 0.00 | 0.77 ± 0.03 | 0.86 ± 0.03 | 0.86 ± 0.02 | 0.73 ± 0.04 | 0.93 ± 0.02 | 0.91 ± 0.01 |
| **Global** | Training | 0.87 ± 0.00 | 0.77 ± 0.01 | 0.92 ± 0.01 | 0.84 ± 0.01 | 0.88 ± 0.01 | 0.81 ± 0.01 | 0.84 ± 0.01 | 0.71 ± 0.01 | 0.87 ± 0.01 | 0.93 ± 0.00 |
|  | Validation | 0.91 ± 0.01 | 0.82 ± 0.05 | 0.93 ± 0.01 | 0.81 ± 0.03 | 0.94 ± 0.02 | 0.82 ± 0.02 | 0.88 ± 0.02 | 0.75 ± 0.03 | 0.89 ± 0.02 | 0.95 ± 0.01 |
|  | Test | 0.84 ± 0.01 | 0.74 ± 0.04 | 0.89 ± 0.01 | 0.79 ± 0.01 | 0.87 ± 0.01 | 0.76 ± 0.02 | 0.81 ± 0.02 | 0.64 ± 0.02 | 0.85 ± 0.03 | 0.91 ± 0.01 |
